# Elaboration of the Homer1 Recognition Landscape Reveals Incomplete Divergence of Paralogous EVH1 Domains

**DOI:** 10.1101/2024.01.23.576863

**Authors:** Avinoam Singer, Alejandra Ramos, Amy E. Keating

## Abstract

Short sequences that mediate interactions with modular binding domains are ubiquitous throughout eukaryotic proteomes. Networks of Short Linear Motifs (SLiMs) and their corresponding binding domains orchestrate many cellular processes, and the low mutational barrier to evolving novel interactions provides a way for biological systems to rapidly sample selectable phenotypes. Mapping SLiM binding specificity and the rules that govern SLiM evolution is fundamental to uncovering the pathways regulated by these networks and developing the tools to manipulate them. We used high-throughput screening of the human proteome to identify sequences that bind to the Enabled/VASP homology 1 (EVH1) domain of the postsynaptic density scaffolding protein Homer1. In doing so, we expanded current understanding of the determinants of Homer EVH1 binding preferences and defined a new motif that can facilitate the discovery of additional Homer-mediated interactions. Interestingly, the Homer1 EVH1 domain preferentially binds to sequences containing an N-terminally overlapping motif that is bound by the paralogous family of Ena/VASP actin polymerases, and many of these sequences can bind to EVH1 domains from both protein families. We provide evidence from orthologous EVH1 domains in pre-metazoan organisms that the overlap in human Ena/VASP and Homer binding preferences corresponds to an incomplete divergence from a common Ena/VASP ancestor. Given this overlap in binding profiles, promiscuous sequences that can be recognized by both families either achieve specificity through extrinsic regulatory strategies or may provide functional benefits via multi-specificity. This may explain why these paralogs incompletely diverged despite the accessibility of further diverged isoforms.

**Impact Statement:** Short linear motifs bind to structurally conserved domains to mediate many protein interactions. Current definitions of motifs, using regular expressions, are over simplified and do not capture what is required for binding. Unbiased proteome interaction screens can refine these definitions. For EVH1 domains considered in this work, screening elucidated Homer1 binding requirements and revealed a residual overlap of specificity of two anciently diverged families, highlighting both opportunities and challenges of multi-specificity in domain-motif interactions.

## Introduction

Biological systems encode numerous modular interaction domains that have expanded into families of paralogs.^1^ Many of these domains recognize stretches of 3-10 sequential residues within intrinsically disordered regions, termed Short Linear Motifs (SLiMs).^2^ Compared to interactions between globular domains, SLiM-mediated interactions are typically weak (µM K_D_) and transient, making them ideal for dynamic processes such as signaling.^3^ SLiMs are also used for protein localization and to coordinate molecular assemblies.^4^

SLiM Binding Domains (SBD) have characteristic motif preferences that are often described using regular expressions that indicate invariant or common residues found in known interaction partners. Matches to these low-information expressions are prevalent throughout the proteome, suggesting that these short sequences alone are poor predictors of binding. Indeed, high-throughput screens of peptide libraries have shown that although most sequences that bind to a given domain share features of a defining motif, not every sequence that matches that motif will bind. Many case studies illustrate how residues outside the core motif can modulate both the affinity and specificity of binding, demonstrating that motifs provide only a partial picture of domain-SLiM molecular recognition.^5,6^

Nature has harnessed SLiM interaction networks to build increasingly complex systems, as evidenced by the rapid expansion of SBD families in higher eukaryotes.^7^ As SBD modules are duplicated, paralogous domains must distinguish their networks to minimize competitive interference and support new functions.^8,9^ Distinct interaction niches can be established in a manner that is extrinsic to the properties of the SBD domain itself, through means such as spatiotemporal compartmentalization of protein expression. Alternatively, this can be achieved intrinsically, at the level of molecular recognition, by changing the sequence to which a domain binds.^10^

The dual promise of unraveling SLiM-mediated networks to uncover novel biology and rewiring such networks to build pathways, e.g., for applications in synthetic biology or as molecular therapies, necessitates a clear understanding of how specificity is established. Here, we explore this question by investigating the divergence of Enabled/VASP homology 1 (EVH1) domains. The human genome encodes 5 EVH1-containing protein families: WASP, SPRED, PP4, Ena/VASP, and Homer.^11,12^ Ena/VASP and Homer are thought to have diverged from a common ancestor, and distinct Ena/VASP and Homer proteins in the choanoflagellate *Salpingoeca Rosetta* suggest that this event occurred at least 600 million years ago.^13,14^

The Ena/VASP family encompasses 3 members: ENAH, VASP, and EVL.^15^ These tetrameric proteins regulate the actin cytoskeleton by serving as a scaffold and acting as processive actin polymerases.^16–18^ Ena/VASP EVH1 domains localize the proteins to sites of actin polymerization, such as filopodia, lamellipodia, and neuronal projections.^19–22^

Like Ena/VASP proteins, the Homer family contains 3 members: Homer1, Homer2, and Homer3.^23^ The constitutively expressed tetrameric Homer proteins function as scaffolds within the post-synaptic density of neurons.^24,25^ Alternatively spliced, monomeric Homer isoforms dynamically regulate signaling pathways by competing with tetrameric Homer and disassembling these complexes in response to neuronal activity.^26,27^ In this capacity, Homer contributes to synaptic plasticity by regulating dendritic spine morphology, perhaps through direct association with actin cytoskeleton proteins such as the actin-bunding protein DBN1.^28,29^ Homer proteins are also expressed in a variety of non-neuronal tissues.^26^ For example, Homer3 negatively regulates T-cell activation by competing with calcineurin for binding to NFATc2.^30^

Sequences of known Ena/VASP partners have revealed the interaction motif [FWYL]PxΦP (Φ = FWYPIALV; x = any amino acid).^31,32^ Several studies have also demonstrated strategies by which regions flanking the core Ena/VASP motif can modulate both affinity and specificity for the EVH1 domain.^32–34^ Fewer Homer binding partners have been validated, and the sequence preferences are less well-defined. Alignments of Homer binding partners suggest a consensus sequence of PPxxF; peripheral regions that modulate the affinity of this motif are poorly understood.^24,31^ Structural alignments of the homologous EVH1 domains in complex with their ligands show that Ena/VASP and Homer families of proteins use partially overlapping binding sites to engage their proline-rich ligands.^35,36^

In this study, we applied high-throughput screening of a library derived from the human proteome to identify Homer1 binding peptides. We found that Homer1 binds to a more general motif than PPxxF and demonstrated how residues flanking the core motif can tune the affinity of this interaction. Notably, compared to the PPxxF motif alone, Homer1 EVH1 domain binds with enhanced affinity to many short segments in the human proteome that match both the Ena/VASP and Homer binding motifs, despite lacking structural features thought to confer these preferences in Ena/VASP proteins. Specifically, we demonstrate that many peptides that contain an N-terminal, overlapping Ena/VASP motif bind with increased affinity to Homer1 while retaining the ability to bind to ENAH. An examination of orthologous EVH1 domains in pre-metazoan organisms suggests that this Homer preference is a vestige of EVH1 domains found in organisms that pre-date the duplication event that gave rise to distinct Ena/VASP and Homer protein families.

## Results

### Structural Comparison of Related EVH1 Domains

Despite only sharing 30% overall sequence identity, the Ena/VASP and Homer EVH1 domain binding sites exhibit a high degree of structural similarity (Figure 1A). Each site is comprised of a proline-binding groove and a neighboring hydrophobic pocket that accommodates the core motif bound by each protein family (Figure 1B).^35,36^

**Figure 1:**
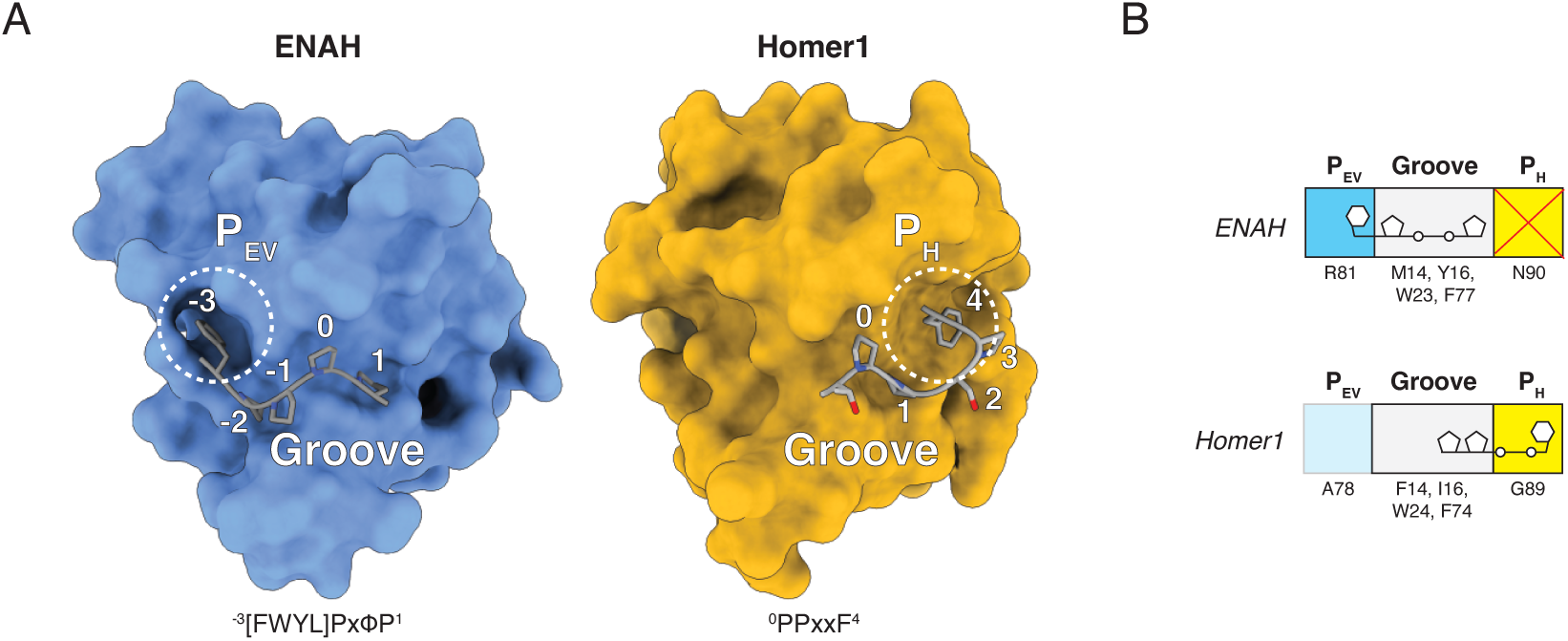
(A) Structures of ENAH (1EVH) and Homer1 (1DDV) in complex with representative ligands. Peptide numbering is relative to the Homer PPxxF motif. For Ena/VASP motifs, the numbering corresponds to the position of the overlapping Pro in both motifs.^35,36^ (B) Cartoon representation of the modular binding pockets along the Ena/VASP and Homer EVH1 domains.

Within the proline-binding grooves of Ena/VASP and Homer, aromatic residues (W23_ENAH_, F77_ENAH_, Y16_ENAH_; W24_Homer1_, F74_Homer1_, F14_Homer1_) accommodate the distinct polyproline-II helical structure adopted by the proline-tracts of bound peptides. We use a unified numbering scheme, defined in Figure 1, in which the Ena/VASP and Homer motifs are indicated as ^-3^[FWYL]PxΦP^1^; ^0^PPxxF^4^. The positioning of aromatic residues within the groove is thought to define the locations of requisite prolines within the preferred motif.^11^ Both families conserve aromatic residues at positions 23 (ENAH)/24 (Homer1) and 77 (ENAH)/77 (Homer1). The unique Ena/VASP aromatic residue, Y16_ENAH_ (I16_Homer1_) engages Pro^-2^, and a preference for Pro at this position is notably absent from the Homer motif. The unique Homer aromatic residue, F14_Homer1_ (M14_ENAH_) engages Pro^1^, which is also engaged by F74_Homer1_ (F77_ENAH_). Thus, F14_Homer1_ seems to reinforce the shared Ena/VASP and Homer preference for proline at position 1.

The peptide-binding specificity of each domain arises from distinct hydrophobic pockets on opposite ends of the proline-binding groove. The Ena/VASP pocket, which we designate here as P_EV_, is primarily formed by K69_ENAH_ (T66_Homer1_), N71_ENAH_ (T68_Homer1_), and R81_ENAH_ (A78_Homer1_), which can form a cation-pi interaction when the residue N-terminal to the motif prolines is aromatic (^-3^**[FWYL]**PxΦP^1^). While the Homer EVH1 domain lacks a well-formed P_EV_, the residues in this region do not sterically occlude this pocket. The Homer pocket, which we call P_H_, is formed by G89_Homer1_, and can accommodate Phe at position 4 (^0^PPxx**F**^4^). In Ena/VASP proteins, this pocket is obstructed by the corresponding N90_ENAH_.

In addition to the core motif, residues in the surrounding sequence can contribute to the affinity and specificity of a peptide for an SBD.^5^ Several affinity-enhancing elements have been discovered for Ena/VASP domains. Acidic residues flanking the ^-3^[FWYL]PxΦP^1^ motif strengthen binding due to positively charged patches along the domain.^32^ C-terminal proline extensions, such as in a peptide from ABI1 (^-3^FPPPP**PPP**^4^), position the motif-trailing residues so they can make favorable hydrophobic contacts near the region corresponding to Homer P_H_.^33^ Finally, Ena/VASP EVH1 domains have been shown to bind a secondary motif at a pseudo-symmetric binding site on the opposite face of the domain (FPPPx_n_**FPPPP**).^33,34^ Few flanking elements have been uncovered for Homer EVH1 domains, although examinations of a handful of known Homer binding partners and mutational studies of a SLiM from the protein DBN1 indicate that Homer prefers an N-terminal Leu (**L**(x)_1-2_PPxxF^4^) and a C-terminal Asp/Asn (^0^PPxxFx**[DN]**).^29^

### A Proteome-wide Screen Elaborates the Homer1 Binding Motif

High-throughput screening of peptide libraries can reveal molecular determinants of SBD interactions.^37^ The T7-pep library encodes the human proteome as 416,611 36-mer peptides and provides a biologically relevant context for identifying putative SBD binding preferences and native interaction partners (Figure 2A).^38^ Hwang et al. formatted this library for bacterial surface display at the C-terminus of the *Escherichia coli* surface protein eCPX and used screening to identify elements that flank the Ena/VASP motif and modulate binding to ENAH.^33,39^

**Figure 2:**
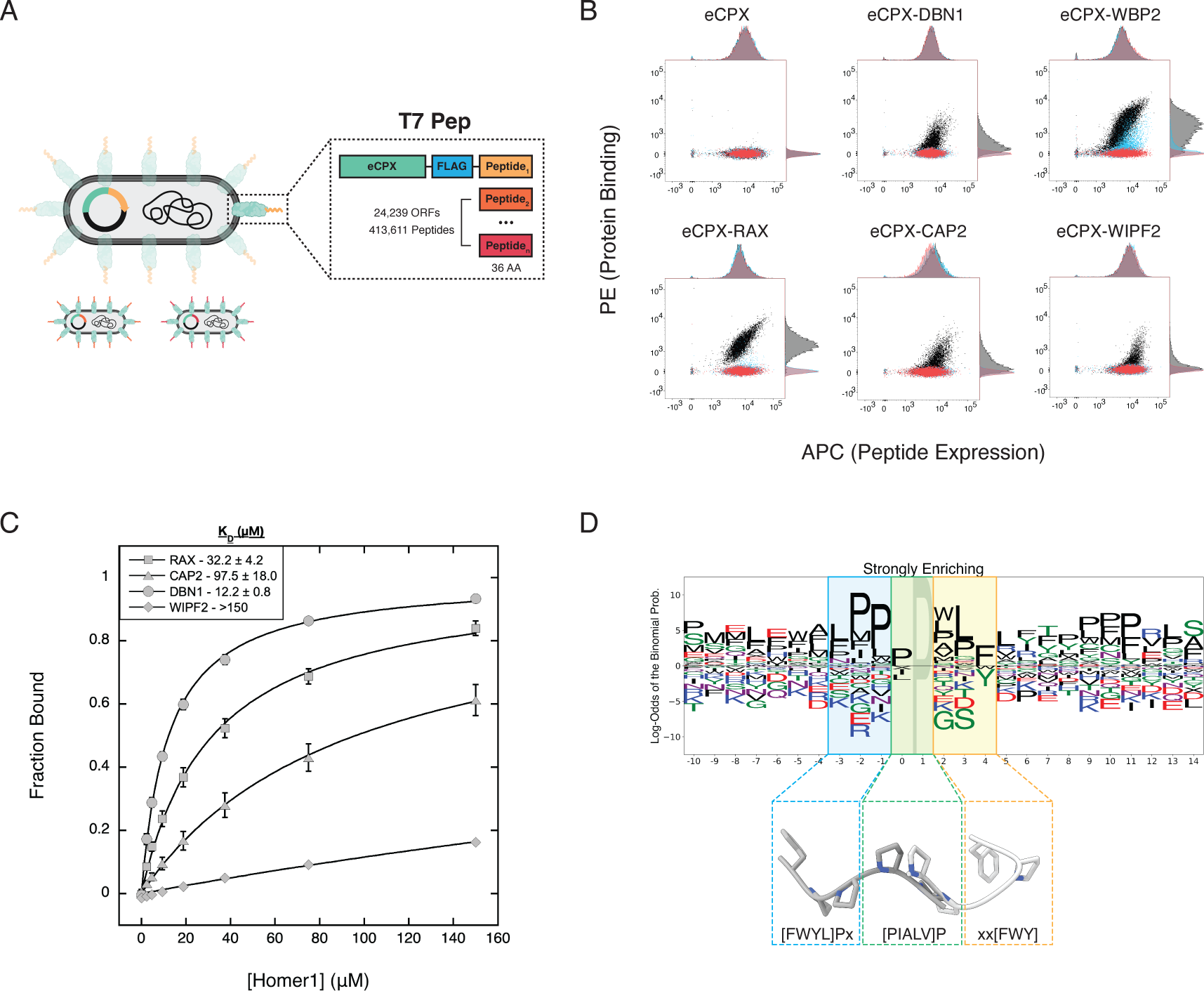
(A) The T7-pep library encodes 36-residue fragments of the human proteome as C-terminal extensions of the *E. coli* surface protein eCPX. (B) Single-clone surface display measurements of control (eCPX, DBN1) and T7-pep derived (WBP2, RAX, CAP2, WIPF2) peptides against Homer1 (black) and binding site mutants (red = Homer1^W24A^, blue = Homer1^G89N^). (C) BLI measurements of control (DBN1) and T7-pep derived (RAX, CAP2, WIPF2) peptides. (D) Sequence logo of peptides strongly enriching for Homer1 binding, aligned by their motif. Structural superposition of ENAH (1EVH) and Homer1 (1DDV) EVH1 domains (not shown) in complex with representative peptides illustrates the similarity between the Homer1 N-terminal motif flank and the Ena/VASP motif (blue = Ena/VASP-like flank, green = overlapping prolines, yellow = Homer-specific motif residues).^35,36^

Preliminary screening of the T7-pep library for peptides that bind the Homer1 EVH1 domain yielded several sequences containing the previously identified PPxxF motif, including peptides from the metabotropic glutamate receptor GRM5, a well-established Homer interaction partner, and WBP2, a protein that has been shown to bind to Homer3 (Table 1).^24,40^ We also discovered matches to this motif in peptides that bind to Homer1 but are not annotated as Homer binding partners, such as the protein RAX, which regulates retinal and neural development.^41^ Tetrameric Homer1 EVH1 domain bound peptides listed in Table 1 on the cell surface, and monomeric EVH1 domain bound to peptide immobilized on a surface via BLI, validating the use of the T7-pep library to identify Homer1 binding sequences (Figure 2B,C).

**Table 1:** T7-pep derived and control (DBN1) peptides that bind to Homer1 via bacterial surface display. The putative Homer1 binding site is shown in bold.

| Name | Sequence |
| --- | --- |
| DBN1 | LNFD~~ELPE~~**PPATF**CDPEEVE |
| WBP2 | **PPPEF**YPGPPMMDGAMGYVQ**PPPPY**PGPMEPPVSG |
| RAX | GFGPPAQSLPAS~~YTPPP~~**PPPPF**LNSPPLGPG~~LQPLA~~ |
| CAP2 | KSSEMNILIPQDGDYREF**P****IPEQF**KTAWDGSKLITE |
| WIPF2 | VEDFP**APEEY**KHFQRIYPSKTNRAARGAPPLPPILR |

In addition to binders with sequences containing the canonical PPxxF motif, we identified several sequences that deviate from this motif by substitution of a small hydrophobic residue at Pro^0^ or substitution of Trp or Tyr at Phe^4^ (Table 2). Testing bacterial clones displaying peptides from the proteins WIPF2 (APEEY) and CAP2 (IPEQF) confirmed binding to Homer1, and these peptides did not bind detectably to established Homer1 mutants with a disrupted proline binding groove (Homer1^W24A^) or with an occluded P_H_ pocket (Homer1^G89N^), suggesting a shared binding mode with the canonical motif (Figure 2B).^35^ BLI measurements showed that these peptides that do not match the consensus motif bound with lower affinity than those including PPxxF sequences (Figure 2B, Table 2). However, low-affinity motifs may be relevant in avid contexts, such as multimeric complexes or proteins containing several motifs.^5^ Notably, full-length CAP2 and WIPF2 contain multiple instances of both canonical and non-canonical Homer motifs.

**Table 2:**
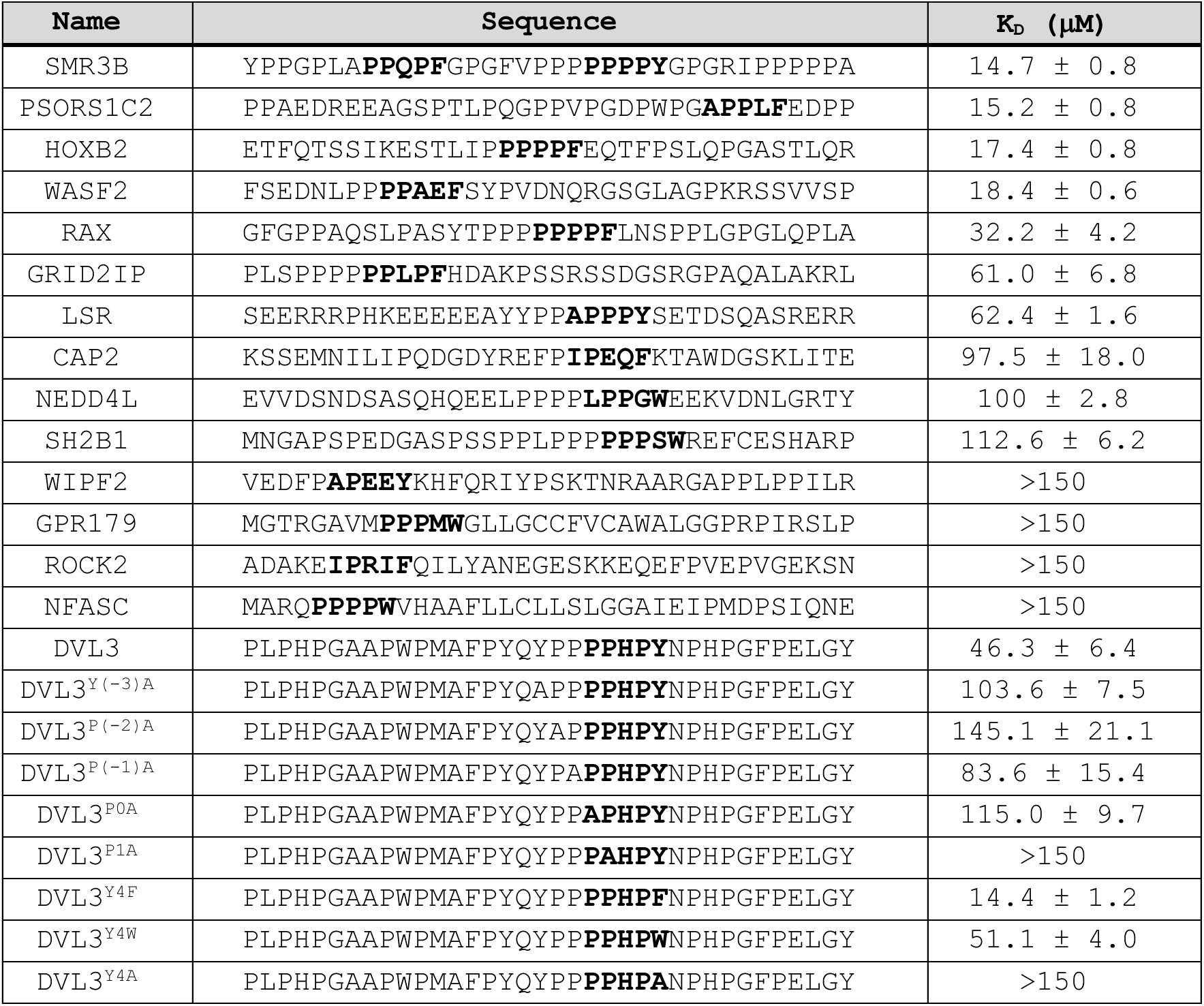
Binding affinities between Homer1 and peptides derived from the T7-pep enrichment screen, measured by BLI. Values represent the average and standard deviation of 3 replicates.

We performed an enrichment screen of the T7-pep library to identify additional Homer1 binders and identified 360 weakly enriching and 62 strongly enriching peptides including the expanded motif ^0^[PIALV]Pxx[FWY]^4^ (Figure S1A, Supplementary Table 1). Subsequent validation using BLI identified peptides that bound with affinities ranging from 15-100 µM, which is within the range of affinities of established Homer binding sequences (Table 2). Interestingly, a peptide from the protein PSORS1C2, which lacks a canonical motif but contains the sequence APPLF, bound with K_D_ = 15.2 ± 0.8 µM, which is similar to the affinity of peptides containing the canonical motif.

### An N-terminal Ena/VASP Motif Enhances Homer1 Binding

Approximately 97% of sequences matching the ^0^[PIALV]Pxx[FWY]^4^ motif that were present in the T7-pep library failed to enrich, even in the avid conditions of the screen. To uncover elements that modulate motif affinity for Homer1, we visualized the screen results using pLogo, which scales residue height according to over- and underrepresentation relative to background amino acid frequencies.^42^ Weakly and strongly enriching sequences were aligned by their motifs and compared to a background of all Homer motif instances in the input library (Figure 2D, Figure S1B).

Notably, the logo of strongly enriching sequences unveiled a preference for the residues ^-3^[LF]PP^-1^ immediately N-terminal to the Homer motif (^-3^**[LF]PP**[PIALV]Pxx[FWY]^4^). These residues resemble the Ena/VASP motif, often encountered as the sequence ^-3^[FWYL]PPPP^1^. We refer to the motif ^-3^[LF]PP[PIALV]Pxx[FWY]^4^ as the extended Homer motif. Comparison of structures of Ena/VASP (PDB: 1EVH) vs. Homer (PDB: 1DDV) EVH1 domains bound to motif-matching peptides show that the two C-terminal prolines of the Ena/VASP motif (^-3^[FWYL]PPPP^1^) bind to Ena/VASP in a pose similar to how the first two prolines of the (^0^PPSTF^4^) motif bind to Homer (Figure 2D).^35,36^ This structural alignment predicts that the N-terminal flanking residues that are enriched in the sequence logo may dock with Homer in a manner similar to the binding of a canonical Ena/VASP ligand to Ena/VASP EVH1, despite the absence of a well-formed P_EV_ pocket or the residue Y16_ENAH_ that which is used by Ena/VASP proteins for Pro^-2^ docking.

Segment polarity protein disheveled homology DVL3, a key player in the Wnt signaling pathway, binds to Homer3 in a yeast two-hybrid assay.^43^ While DVL3 does not contain a canonical PPxxF motif, a peptide containing the non-canonical motif and an Ena/VASP-like N-terminal flank (^-3^YPPPPHPY^4^) enriched in our screen and bound with K_D_ = 46.3 ± 6.4 µM. Alanine substitutions throughout this sequence were tolerated to varying degrees (Table 2). Within the core motif, the substitution of Pro^1^ or Tyr^4^ with Ala resulted in peptides without appreciable binding to Homer1, while substitutions of Pro^0^ caused nearly a 3-fold reduction in binding affinity (Figure 3A). Mutations within the Ena/VASP-like flank had a similar impact on overall binding affinity as substitution of Pro^0^ or the aromatic residue at position 4, highlighting the extent to which motif-external elements can influence binding. Interestingly, most of the BLI-validated sequences contained at least a partial match to the Ena/VASP-like flank, which was absent from non-binding sequences with non-canonical motifs. These results demonstrate that an Ena/VASP-like flank can enhance Homer1 binding and compensate for a suboptimal core Homer motif.

**Figure 3:**
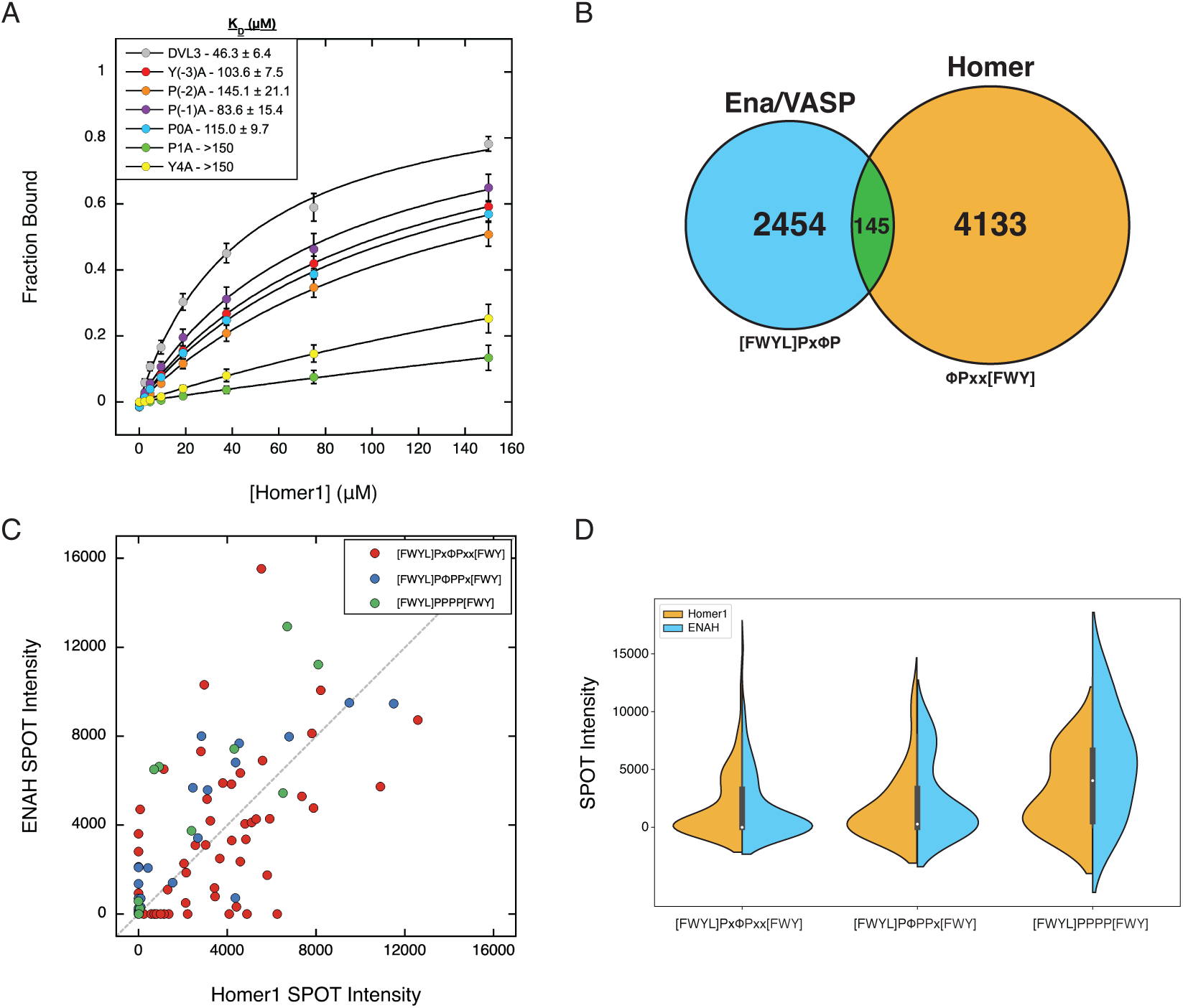
(A) Alanine mutations throughout the Homer1 motif and N-terminal Ena/VASP-like flank in DVL3 diminish binding to Homer1. (B) Number of peptides that match the Ena/VASP (blue), Homer (gold), and overlapping motifs (green; ^-3^[FWYL]PxΦPxx[FWY]^4^, ^-2^[FWYL]PΦPPx[FWY]^4^, or ^-1^[FWYL]PPPP[FWY]^4^) within disordered regions of the human proteome. (C,D) SPOT intensities for ENAH and Homer1 binding to human peptides containing overlapping Ena/VASP and Homer motifs.

### The Homer1 and ENAH EVH1 Domains have Incompletely Diverged

The identification of an N-terminal Ena/VASP motif as a Homer affinity-enhancing flanking element is perplexing. Homer proteins are highly expressed in neurons, where Ena/VASP proteins play important roles in neuronal migration and axon guidance.^22^ If Ena/VASP EVH1 domains are similarly capable of binding these overlapping motifs, that would imply the potential for competitive binding between the two protein families that have different functions.

The intrinsically disordered regions (IUPRed > 0.45) of the human proteome contain 103 instances of a Homer motif flanked by a potentially affinity-enhancing Ena/VASP-like sequence, ^-3^**[FWYL]Px**ΦPxx[FWY]^4^ (Figure 3B, Supplementary Table 2).^44^ To determine whether peptides that match the core motifs of both domains bind to both Ena/VASP and Homer EVH1 domains, we tested the interaction of these peptides with tetrameric EVH1 constructs using a peptide SPOT array (Figure 3C, Figure S2). Many peptides bound to both domains and the distributions of intensities for all peptides were remarkably similar for Homer1 and ENAH (Figure 3D). BLI measurements confirmed that many peptides bind to both ENAH and Homer1 with similar affinities (Table 3). A further 42 Homer motif-matching peptides contain an N-terminally overlapping Ena/VASP motif in a frame that does not structurally align to the Ena/VASP binding mode, that is, a match to ^-2^**[FWYL]P**ΦPPx[FWY]^4^ or ^-1^**[FWYL]**PPPP[FWY]^4^ (Figure 3B). While many peptides within these classes were similarly capable of binding Homer1, these peptides showed a slight bias toward binding ENAH based on their distribution of SPOT intensities (Figure 3C,D).

**Table 3:**
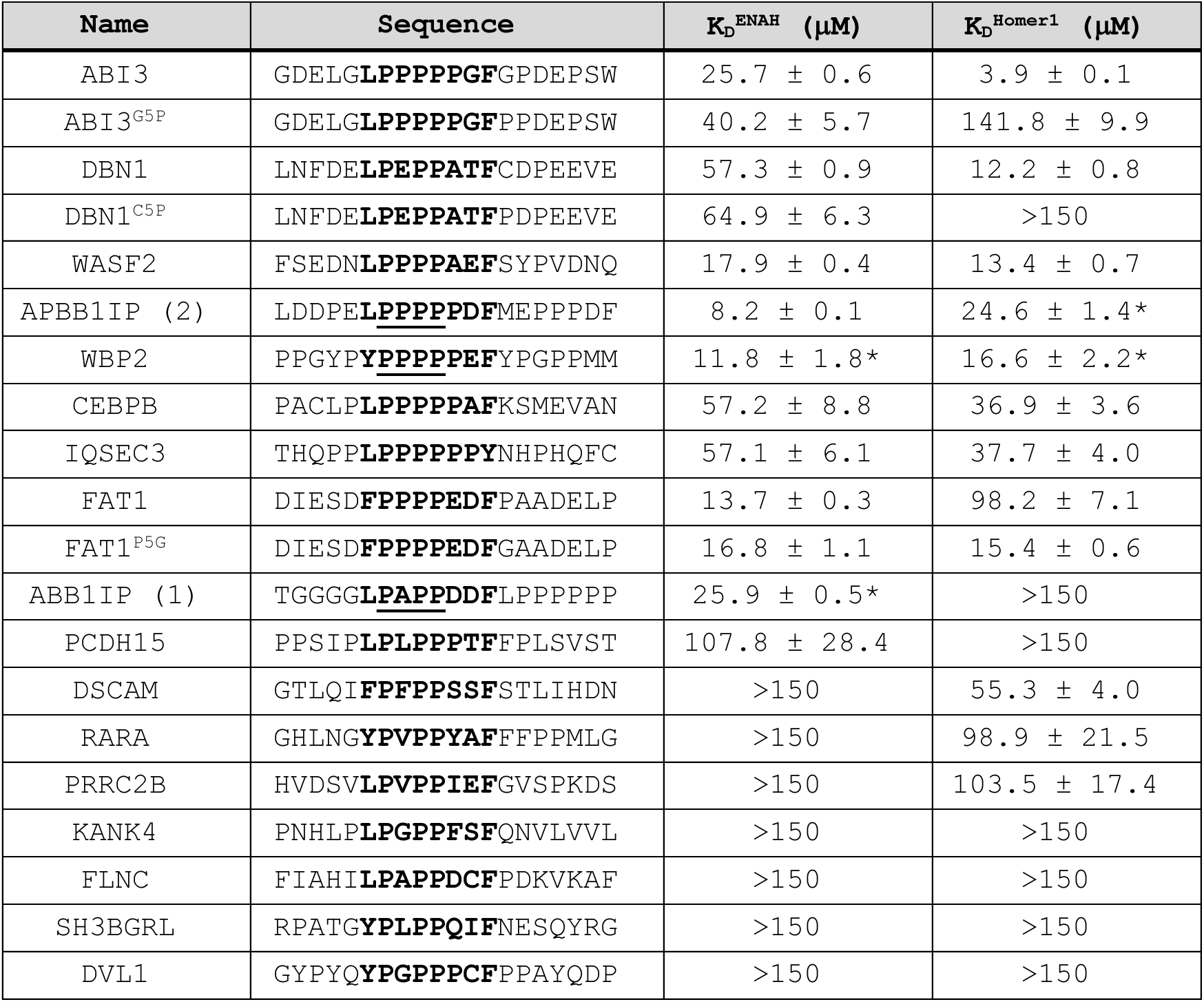
Binding affinities of bioinformatically-identified peptides containing the extended Homer motif for Homer1 or ENAH, determined using BLI. Measurements marked with an asterisk (*) are peptides for which SPOT array binding was maintained even when the motif was disrupted via the mutation of the underlined residues to glycine. Values represent the average and standard deviation of 3 replicates.

### Single Motif-Flanking Residue Modulates Binding to Homer1

Of the 140 peptides that contain overlapping Ena/VASP and Homer motifs that we tested, 58 bound detectably to Homer1 and 55 to ENAH. Most peptides were not highly specific for either domain, with 46 peptides binding to both. Several overlapping peptides exhibited modest selectivity in the SPOT array assay, such as FAT1 (FPPPPEDF) and ASPP2 (YPPPPY), which bound to ENAH 2 to 3-fold better than Homer1. FAT1 also demonstrated a near 10-fold preference for ENAH (K_D_ = 13.7 ± 0.3 µM) over Homer1 (K_D_ = 98.2 ± 7.1 µM) via BLI, corroborating the differences in SPOT intensity (Table 3). The weak affinity of Homer1 for FAT1 is surprising given the presence of both a canonical PPxxF motif and an Ena/VASP-like flank, and we hypothesized that features in the motif-surrounding sequence may disfavor the FAT1-Homer1 interaction.

To identify potential negative design elements, we created logos for the sequences that bound specifically to Homer1 or ENAH by SPOT array. This revealed that Homer1, but not ENAH, disfavored Pro at position 5, immediately C-terminal to the core Homer motif (Figure 4A). Notably, both FAT1 and ASPP2 encode a Pro in this position. The crystal structure of Homer2 bound to a peptide from DBN1 (PDB: 5ZZ9) suggests that this residue, when not Pro, can hydrogen bond with the residue in position 3 using its backbone amide hydrogen (Figure 4B).^29^ This interaction may orient the aromatic residue in position 4 for docking within P_H_, explaining the specificity of this effect for Homer1. Indeed, removing this element in FAT1 (P5G) yielded a peptide that bound six-fold tighter to Homer1, but this mutation had almost no effect on ENAH binding (Figure 4C). Similarly, introducing Pro at this position in peptides from ABI3 (G5P) and DBN1 (C5P) substantially decreased the affinity for Homer1 with minimal impact on ENAH binding. The role of proline at position 5 is an elegant example of a structural strategy that can increase the binding specificity of peptide segments that match both the Ena/VASP and Homer binding motifs.

**Figure 4:**
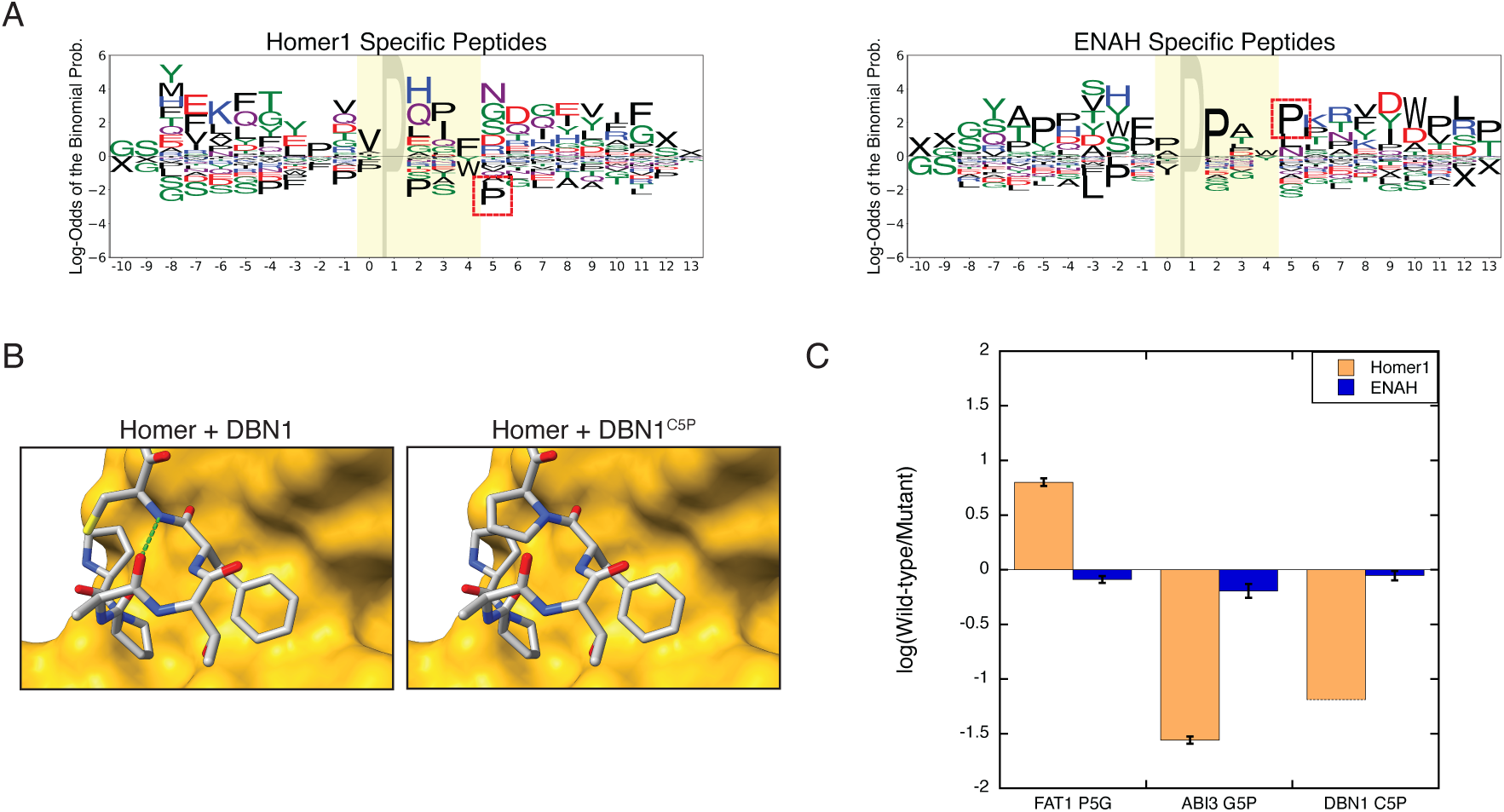
(A) Logos for bioinformatically-identified peptides containing the extended Homer motif that preferentially bind to Homer1 or ENAH by SPOT array. Sequences are aligned by their Homer1 motif. The red box indicates the position of the Pro^5^ Homer1 negative element. (B) Homer2 EVH1 domain bound to a peptide from DBN1 demonstrates that the residue in position 5, when not Pro, can hydrogen bond (green) with the residue in position 3 (5ZZ9).^29^ (C) The fold- change in binding affinity of ENAH and Homer1 EVH1 domains for peptides in which a Pro has been removed from (FAT1^P5G^) or introduced (ABI3^G5P^, DBN1^C5P^) in the position following the Homer1 motif.

### Transformation of EVH1 Binding Preferences

Co-crystal structures of Ena/VASP and Homer EVH1 domains bound to representative ligands have revealed residues within the binding sites that, through their contacts, appear crucial for defining the reported binding motifs. To identify the extent to which this small set of residues is responsible for conferring the motif preferences, we attempted to convert the binding preferences of Homer1 and ENAH by swapping four binding site residues in ENAH, representing 5% of the differences between these domains (ENAH^Homer1-like^; M14F/Y16I/R81A/N90G) (Figure 5A). As demonstrated by a single-substitution SPOT array of the promiscuous peptide from ABI3, ENAH was most sensitive to substitutions within the Ena/VASP motif (^-3^LPPPP^1^), whereas Homer1 was sensitive to substitutions within the Homer motif (^0^PPPGF^4^) (Figure 5B). The tolerance for deviations from the canonical motifs for each domain and the inability to resolve the expected contribution of N-terminal residues to Homer1 binding is likely due to the use of a high-affinity ligand in a multivalent context. Notably, ENAH^Homer1-like^ exhibited Homer1-like binding properties, both in terms of the decreased sensitivity to substitutions in the Ena/VASP motif and the acquired requirement for a large hydrophobic residue at position 4 (Figure 5D). Unlike Homer1, ENAH^Homer1-like^ had a strong preference for valine at position 4, implying that additional differences between these domains are responsible for fine-tuning the motif. Nevertheless, the ability to recapitulate most Homer1 preferences within a markedly different background verifies that just a few residues play a pivotal role in defining EVH1 domain specificity.

**Figure 5:**
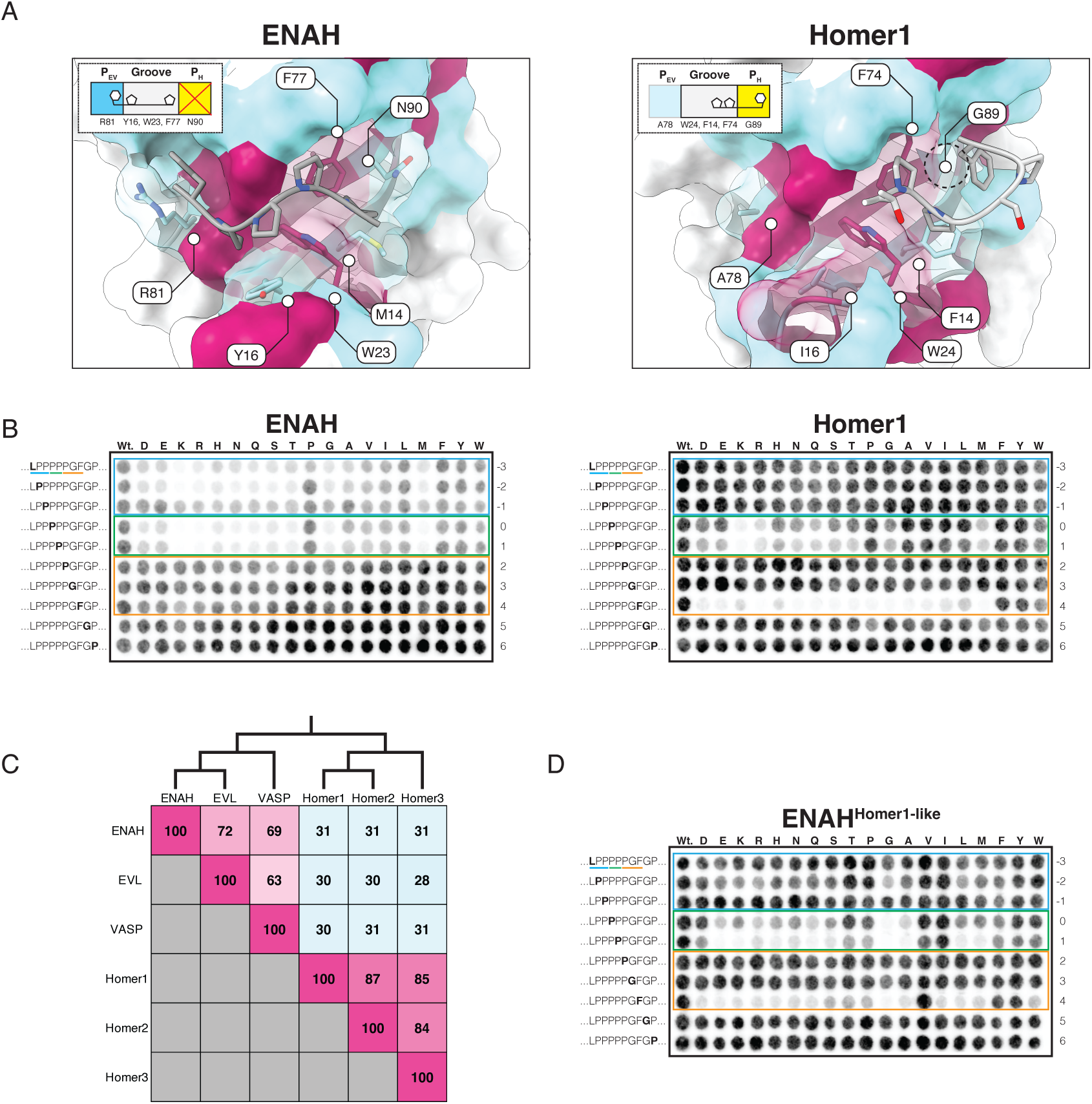
(A) ENAH (EVH1) and Homer1 (1DDV) in complex with their representative ligands. Residues within 7Å of the binding site are colored by whether they are shared (maroon) or different (cyan) between these paralogous domains.^35,36^ (B) Mutational analysis of the overlapping motif in ABI3 binding to ENAH and Homer1 EVH1 domains. In each row, the substituted residue is marked in bold text. (C) Percentage identity matrix of the human Ena/VASP and Homer family EVH1 domains. The tree corresponds to the pairwise sequence identity between domains. (D) Mutational analysis of the overlapping motif in ABI3 binding to the ENAH^Homer1-like^ EVH1 domain.

### An Ortholog from *Dictyostelium discoideum* Sheds Light on the Evolution of EVH1 Binding Preferences

Ena/VASP and Homer are highly conserved in metazoans, and both families are present in choanoflagellates, the single-cell organisms most closely related to metazoans.^14^ Several phylogenetically remote amoebas lack Homer proteins but encode a singular Ena/VASP protein, as defined by the presence of both EVH1 and EVH2 domains. We hypothesized that these orthologs, seemingly from organisms that pre-date the gene duplication event that gave rise to Homer, may offer insight into the evolution of binding specificity within this paralogous family.

In the soil-dwelling amoeba *Dictyostelium discoideum*, the Ena/VASP protein, ddVASP, is required for filopodia formation, chemotaxis, and macroendocytosis.^45,46^ As assessed by sequence identity, the *D. discoideum* EVH1 domain (which we refer to here as ddEVH1) is similarly diverged from the human Ena/VASP (35%) and Homer (31%) EVH1 domains (Figure 6A). An AlphaFold2 model of ddEVH1 reveals that the region corresponding to the peptide binding site in Ena/VASP and Homer exhibits a blend of features from both human protein families (Figure 6B).^47,48^ Like Homer, but not Ena/VASP, ddEVH1 lacks a well-formed P_EV_ (T80_ddEVH1_ and A78_Homer1_ vs. R81_ENAH_). Like Ena/VASP, but not Homer, ddEVH1 has an occluded P_H_ (N89_ddEVH1_ and N90_ENAH_ vs. G89_Homer1_). Additionally, whereas Ena/VASP and Homer EVH1 domains each contain three aromatic residues in the proline binding groove, ddEVH1 contains a tetrad that includes both the unique Ena/VASP (Y16_ddEVH1_ and Y16_ENAH_ vs. I16_Homer1_) and Homer (F14_ddEVH1_ and F14_Homer1_ vs. M14_ENAH_) aromatic residues. Sequence analysis of the Ena/VASP proteins from other amoebas demonstrates that this binding site architecture is highly conserved (Figure S4).

**Figure 6:**
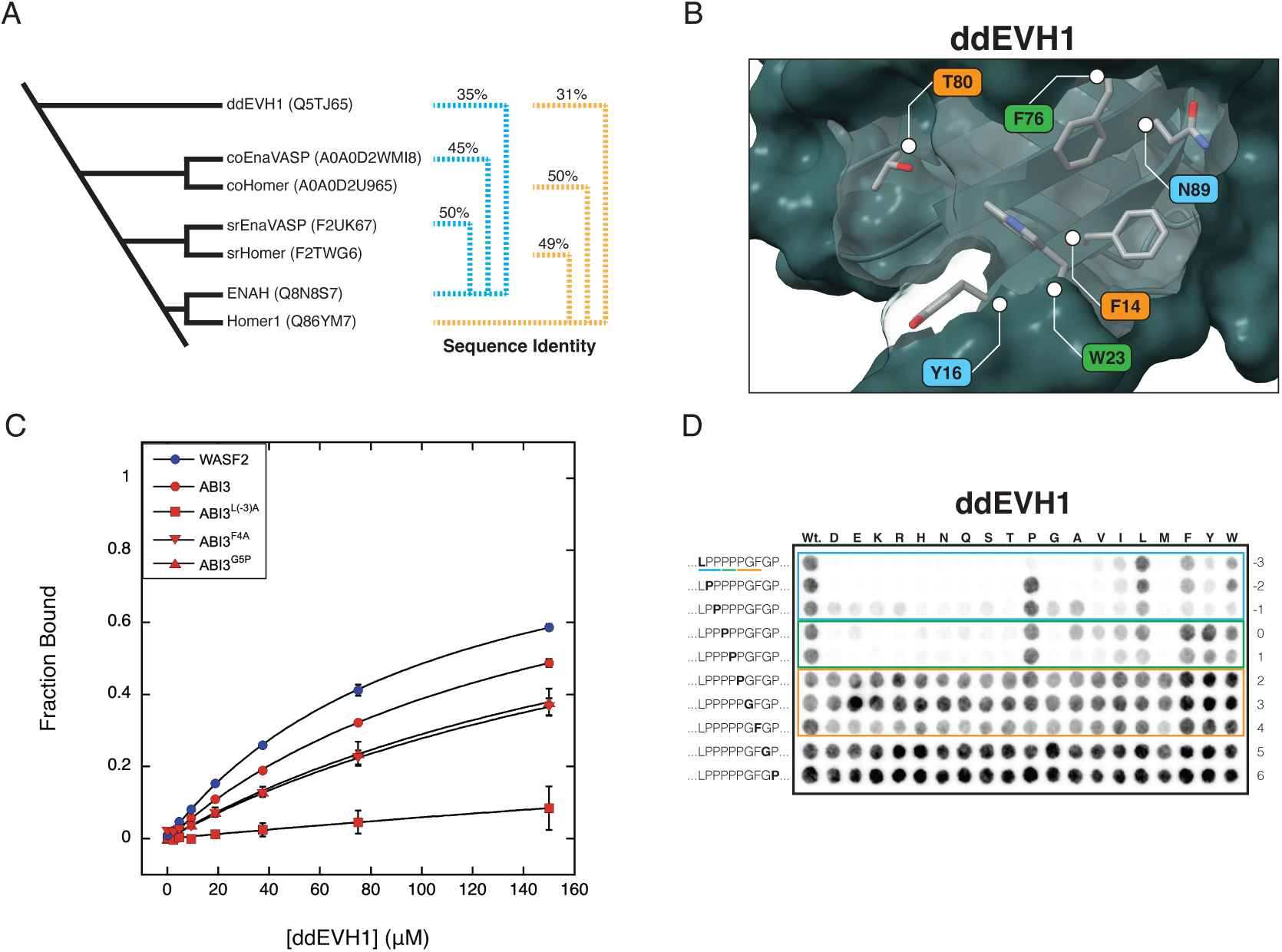
(A) Dendrogram representing the positioning of diverse organisms containing Ena/VASP and Homer proteins along the tree of life. The sequence identity for Homer EVH1 domains (and ddEVH1) is shown relative to human Homer1 (yellow), and the sequence identity for Ena/VASP EVH1 domains is shown relative to human ENAH (blue). (B) AlphaFold2 structural model of ddEVH1 (Q5TJ65). The color of each label reflects whether the residue is similar to the corresponding residue in Homer (yellow), Ena/VASP (blue), or both (green). (C) BLI measurements for ddEVH1 and promiscuous peptides from WASF2 and ABI3. (D) Mutational analysis of the overlapping motif in ABI3. In each row, the substituted residue is marked in bold text.

We tested ddEVH1 for binding to several human peptides identified in this study (Figure 6C). Like ENAH and Homer1, ddEVH1 bound to peptides from ABI3 and WASF2, though the affinity for each was weak (K_D_ > 100 µM). Alanine substitution of the N-terminal Leu^-3^ (^-3^**L**PPPPPGF^-4^) in ABI3 (ABI3^L(-3)A^) abrogated ddEVH1 binding, while substitution of the C-terminal Phe^4^ (ABI3^F4A^), or replacement of the motif-trailing Gly^5^ with Pro (ABI3^G5P^), had modest effects on binding. In these ways, the ddEVH1 motif is similar to the human Ena/VASP motif, despite, like Homer1, containing differences in key binding site residues. Nevertheless, ddEVH1 bound poorly to the ENAH-binding peptides from DBN1 and FAT1 (data not shown).

Single-substitution SPOT array of ABI3 demonstrated that, in contrast to ENAH, ddEVH1 strongly preferred Leu over an aromatic residue at position −3 (Figure 6D). This is likely caused by the absence of cation-pi interactions formed in ENAH by R81_ENAH_ and may explain the weak binding of ddEVH1 to FAT1 (^-3^**F**PPPPEDF^4^). The array also revealed that ddEVH1 has a stronger preference than ENAH for proline residues at positions −2 through 1. This may indicate that the proline binding groove, with its extra aromatic residue, compensates for a poorly formed P_EV_ pocket. DBN1, which does not bind to ddEVH1, lacks a proline at the −1 position (^-3^LP**E**PPATF^4^). Lastly, ddEVH1 exhibits a weak sensitivity to the replacement of Phe^4^ with a non-aromatic residue, which aligns with our finding that alanine substitution at this position (ABI3^F4A^) had a modest effect on binding, as measured by BLI. This preference is surprising, given the absence of P_H_ that accommodates the binding of this residue to Homer EVH1 domains.

Our biochemical characterization of ddEVH1 suggests a preference for the motif [LF]PPPP, with less tolerance for deviation within this sequence than is observed in human Ena/VASP orthologs. 113 *D. discoideum* proteins contain an [FL]PPPP motif. Of these, 22 have human orthologs, with several being annotated as participating in actin regulation or neuronal processes (Supplementary Table 3). Furthermore, the ABI protein in *D. discoideum*, a known binding partner of ddEVH1, contains 6 copies of the sequence [LF]PPPP (UniProt ID: Q55FT9).^49^ This fact, together with our observation that peptides matching this motif bind ddEVH1 an order of magnitude weaker than ENAH and Homer1, suggests that multivalency may be crucial for obtaining physiologically relevant binding affinities in amoeba Ena/VASP systems.

### A Homer Ortholog from *Capsaspora owczarzaki* can Discriminate Against ENAH Motifs

The incomplete divergence of the Ena/VASP and Homer binding profiles may imply that evolutionary paths to distinct interactions are long (requiring many mutations) or rare (requiring very specific combinations of mutations), such that fully differentiated binding properties are unlikely to arise even over long time periods. This could be the case, for example, if most binding-site mutations that lead to diverged binding compromise the structural integrity of the domain. The advent of multicellularity provided a new mechanism by which biological systems can spatiotemporally restrict the co-localization of proteins, segregating proteins into distinct niches without the need for divergence at the level of molecular recognition. Given the potential for deleterious competition between Ena/VASP and Homer, we hypothesized that pre-metazoan unicellular organisms, may face greater pressure to diverge their interface residues.

Our search for orthologs in single-cell species identified both an Ena/VASP and Homer protein in the eukaryote *Capsaspora owczarzaki*, a member of the Filasteria clade, the sister group to choanoflagellates and metazoans (Figure 6A). The EVH1 domain in the Homer protein, which we call coHomer, shares 50% sequence identity to human Homer1. While the architecture of the proline groove and P_H_ are nearly identical to Homer1, coHomer uniquely incorporates an acidic residue (E78) within the P_EV_ pocket (Figure 7A). This mutation may disfavor the engagement of peptides that bind well to Ena/VASP by disrupting the docking of a hydrophobic residue in P_EV_ or by repelling peptides containing acidic residues often found upstream of the motif in Ena/VASP binders.^32^

**Figure 7:**
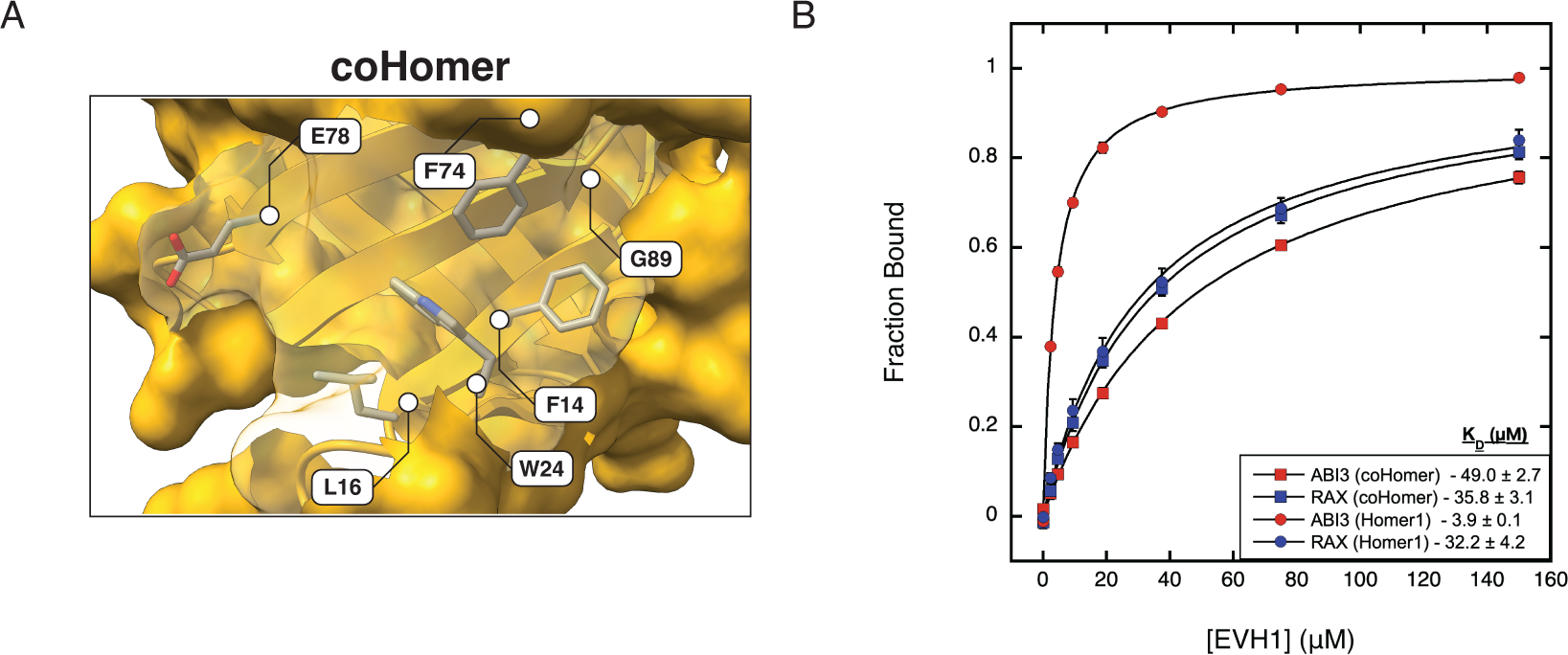
(A) AlphaFold2 structural model of coHomer (UniProt: A0A0D2U965). (B) BLI binding curves comparing Homer1 (circle) and coHomer (square) binding to peptides from ABI3 and RAX (sequences in Tables 1 & 2).

We compared the binding of multiple peptides to both Homer1 and coHomer EVH1 domains using BLI (Figure 7B). Peptides that included acidic residues within a 5-residue window N-terminal to the motif (ABI3, DBN1, and FAT1^P5G^) bound to coHomer with at least a 10-fold reduction in affinity compared to Homer1 (Table 4). DVL3, which contains an overlapping motif but lacks N-terminal acidic residues, also exhibited a reduction in binding affinity. RAX, which lacks the Ena/VASP P_EV_ docking residue (^-3^**[FWYL]**PxΦP^1^), bound equivalently to both Homer1 and coHomer. This trend suggests that coHomer disfavors binding to Ena/VASP-binding peptides largely by repelling N-terminal acidic residues, but also through placing a charge in P_EV_. Introducing this glutamate into the Homer1 EVH1 domain (Homer1^A78E^) alone was insufficient to evoke a similar change in affinity (Figure S5A). This may be due to the greater flexibility of Glu in Homer1 compared to the corresponding residue in coHomer, which is packed against L85 and V80 (Figure S5B). While multiple mutations may be required to impart the binding preferences of coHomer, our results demonstrate that further diverged molecular recognition profiles of Homer and Ena/VASP are evolutionary accessible but have not been selected for in metazoan lineages.

**Table 4:**
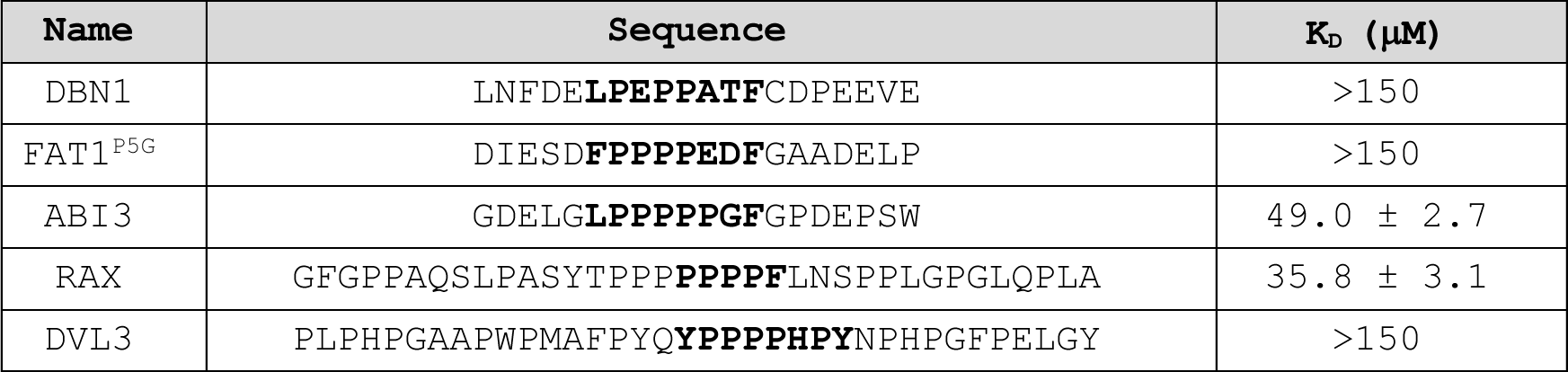
coHomer binding affinities measured by BLI. Values represent the average and standard deviation of 3 replicates.

## Discussion

### Homer1 Binding Preferences

Homer, a constitutive tetramer, carries out its biological function by crosslinking (or, in the case of immediate early monomeric Homer isoforms, disassembling) complexes of partners using its EVH1 domain.^23^ Identifying these binding partners is key to understanding the biological role of Homer proteins. Because of the relatively weak affinity (low- to midmicromolar) of EVH1-mediated binding and the dynamic nature of the postsynaptic density, many of these interactions are likely transient and only populated in response to specific stimuli. For this reason, Homer-mediated interactions may evade standard techniques for identifying native protein-protein interactions, and this may explain the low recovery of annotated binding partners in various Homer co-IP mass spectrometry studies.^50,51^ Limitations of these experiments also stem from their tendency to focus on a single tissue type, despite the wider expression of Homer.^26^

Most Homer binding partners have been identified in low-throughput, candidate-driven studies by virtue of the inclusion of a canonical PPxxF motif, first identified in metabotropic glutamate receptors.^24,28^ This led to the circular reinforcement of PPxxF as the Homer binding motif, potentially to the exclusion of partners that lack this sequence. Several deviations from this motif have been noted in the literature, but this has not led to a broader exploration of Homer binding preferences.^52,53^

High-throughput screening of peptide libraries is a powerful technique for mapping domain binding landscapes in an unbiased way.^54^ Proteome-derived libraries such as Pro-PD and T7-pep allow for the identification of candidate binding partners of potential biological relevance.^33,37,38^ Even for library hits that do not interact in a cell, biochemical binders nonetheless offer insight into the diverse ways that a domain can engage its ligands.

In this study, we screened T7-pep to identify Homer1 binders and uncovered features that support binding. Our results demonstrate that the reported core Homer motif of PPxxF is overly restrictive and that sequences matching a more general motif of [PIALV]Pxx[FWY] can bind to the Homer1 EVH1 domain. Additionally, we identified specific examples of how the sequence context flanking the core motif, such as a leading Ena/VASP motif-like segment or a trailing proline, can modulate the affinity and specificity of this interaction. These findings build on and contextualize the previous discoveries that many Homer ligands contain an N-terminal, affinity-enhancing Leu, and that mutation of an N-terminal FPP diminishes the binding of Homer3 to FAT1.^29,55^ By combining our findings and reported Homer preferences from the literature, we define an updated Homer motif [fwyl]pp[PIALV]P..[FWY][^p][dn], where uppercase letters reflect obligate positions and lowercase letters indicate positive and negative elements that can alter the affinity of this interaction.^29,56^

### The Evolution of Binding Preferences in EVH1 Domains

Homer and Ena/VASP EVH1 domains share a common ancestor, and these distinct families are present in the pre-metazoan *Salpingoeca rosetta*, indicating that the family-expanding duplication event predated the emergence of metazoans some 600 million years ago (Figure 6A).^13^ Many amoebas, further remote from metazoans along the tree of life, encode a singular Ena/VASP protein but lack Homer orthologs. The absence of Homer further back in the tree suggests that these organisms predate the duplication event that gave rise to Homer. Thus, we hypothesized that EVH1 domains from amoebas may represent the ancestral domain’s binding preferences.

The EVH1 domains from amoebas share near-equal sequence identity with domains from both human Ena/VASP and Homer. Furthermore, the binding site architecture of these domains blends features from both human families. While we cannot rule out that the binding preferences of these EVH1 domains may have drifted over time, proteins at the center of interaction networks are buffered against drift due to the extensive co-evolution required to maintain their interactomes.^57–60^ In support of this, even those EVH1 domains from amoebas with low sequence identity share the binding site residues we identified as key for defining motif preferences, suggesting the persistence of a common binding mode (Figure S4). Together, these observations suggest a model whereby subfunctionalization of an ancestral binding site similar to that in ddEVH1 birthed distinct Ena/VASP and Homer families.

In support of this model, we observed that ddEVH1 binds weakly to the motif [FL]PPPP and exhibits a slight preference for the C-terminal aromatic residue found in the Homer motif. Seemingly, selective reinforcement of pre-existing preferences for hydrophobic residues flanking the prolines, as opposed to *de novo* emergence, established the higher-affinity motifs that are recognized by metazoan EVH1 domains (Figure 8). As evidenced by the affinity of Homer1 for sequences containing an Ena/VASP-like N-terminal flank, Homer did not lose its preference for the ancestral motif, which exists independent of a well-formed P_EV_. Similarly, despite lacking a distinct P_H_, the ENAH EVH1 domain can utilize this site to bind to a motif-trailing proline in ABI1 (^-3^FPPPPP**P**P^4^).^33^ Furthermore, negative design elements did not arise to eliminate binding overlap between the human Ena/VASP and Homer families.

**Figure 8:**
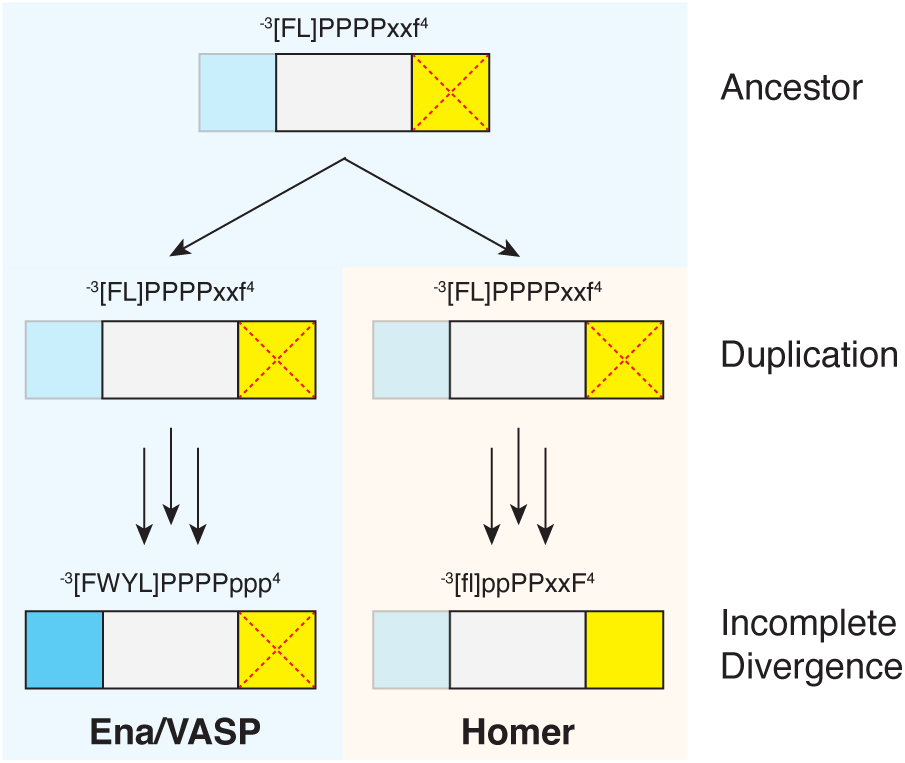
An evolutionary model of EVH1 divergence. Uppercase letters reflect obligate positions and lowercase letters indicate positive elements that can alter the affinity of this interaction The EVH1 ancestor of Ena/VASP and Homer likely bound to a motif akin to that preferred by ddEVH1, with a poorly formed PEV (transparent) and occluded PH (red cross mark). Incomplete subfunctionalization of the core binding site resulted in a Homer EVH1 domain that retained its preference for an Ena/VASP motif.

Ena/VASP and Homer EVH1 domains may not have diverged further because changes that enhance selectivity compromise binding. However, the sole Homer protein in the premetazoan *Capsaspora owczarzaki* robustly discriminates against sequences containing an Ena/VASP motif and affinity-enhancing acidic residues, demonstrating that further diverged binding profiles are accessible. Why then did the Homer preference for the Ena/VASP motif persist in metazoan lineages? We posit that the retention of this preference in Homer is either functionally beneficial or neutral with respect to fitness but preserved due to constraints on drift, for example, arising from high interaction connectivity.

### Rationalizing Overlap in the Ena/VASP and Homer Recognition Landscape

SLiMs play pivotal roles in cellular physiology, and mutations that disrupt these sequences can reduce fitness and lead to human disease.^4,61^ Fitness can also be compromised by competition between proteins for binding, leading paralogs to develop specificity niches, either by diverging their molecular recognition profiles or by spatiotemporal compartmentalization.^62–64^ Yeast SH3 domains provide examples of both strategies.^65,66^ A peptide from the protein Pbs2 binds to the Sho1 SH3 domain with high specificity, indicating that specificity can be encoded in molecular recognition.^67^ On the other hand, a peptide from Pex14 binds promiscuously to a number of SH3 domains, but it is sequestered with its cognate SH3 domain (Pex13) in the peroxisome. Similarly, the Ena/VASP and Homer EVH1 domains have diverged their recognition profiles sufficiently for specific, non-promiscuous ligands to exist. Perhaps like the Pex14-Pex13 interaction, promiscuous peptides that bind to both Ena/VASP and Homer are spatiotemporally restricted from encountering both simultaneously.

Alternatively, these peptides may convey a beneficial function. The intricate landscape of cellular processes is dependent on conditional interactions, and one way of achieving this type of regulation is through direct competition for binding sites, such as that between Homer3 and calcineurin for binding to NFATc2.^30,68,69^ Competitive binding between constitutively expressed tetrameric Homer and the immediate early monomeric isoform is central to Homer function.^26,27^ Competitive interactions with Ena/VASP proteins may play a similar role for motifs that bind to both families.

Several proteins that contain dual-binding peptides stand out as candidate functional partners for both Ena/VASP and Homer based on localization, interaction, and GO term data (Figure S3). Among these are ABI3 and WASF2, two members of the pentameric WAVE complex that regulates Arp2/3 mediated actin polymerization.^70^ The WAVE complex interacts directly with Ena/VASP proteins to drive enhanced actin assembly, and this association is conserved as far back as *D. discoideum*.^49,71^

Given the role of Homer in synaptic plasticity - an actin remodeling event - a direct connection between Homer and actin polymerization machinery is conceivable. Circumstantial evidence suggests that Homer proteins may also interact with the WAVE complex. In neurons, the WAVE complex plays a crucial role in orchestrating the development of diverse actin-filament protrusions.^72^ ABI3 has been shown to regulate dendritic spine morphology and synapse formation, and the peptide from ABI3 is the tightest Homer1 binder identified in this study (K_D_ = 3.9 ± 0.1 µM).^73^ Through its interaction with various proteins, Homer can influence both the emergence and maturation of dendritic spines.^74,75^ Homer3 knockdown in neutrophils impairs the recruitment of the WAVE2 complex to the leading edge of motile cells.^76^ In the cerebral cortex of mice, Homer1 dissociates from a complex containing the Arp2/3 members ARPC2, 4, and 5 in an activity-dependent manner.^50^

The small number of interface residues that constitute SLiM-mediated interactions makes them highly amenable to regulation by post-translational modification. Motif phosphorylation is a well-documented switch for SLiM binding (e.g phosphorylation of the NPxY motif in the integrin *β*3 subunit shifts binding from talin to Dok1).^68,77^ Phosphorylation can also modulate the affinity of a motif, and phosphorylation of the non-interface Ser and Thr residues in the mGlur5 Homer binding peptide (TPPSPF) by Cdk5 increases affinity for Homer1.^78^

Our study uncovered that Homer1 can bind to sequences containing Tyr^4^ within the core motif, but is intolerant to acidic residues and presumably pTyr^4^ at this position. However, in the context of an overlapping motif, this modification may enhance binding to Ena/VASP, given the preference for motif-flanking negative charges.^32^ Of the 46 dual-binding peptides identified in this study, 20 contain Tyr^4^ and 5 of these residues are annotated as phosphorylation sites.^79^ One of these proteins, the non-receptor kinase Activated CDC42 kinase 1 (TNK2) has a documented role in neurite migration, and phosphorylation of this Tyr plays a role in glioma oncogenesis.^80,81^

In summary, our results suggest that the Homer interaction landscape remains only partially explored, and we present new motif definitions and candidate binding partners to facilitate the discovery of novel interactions. Furthermore, these findings provide a case study into the evolution of molecular specificity in SLiM binding domains by documenting the incomplete divergence within this paralogous family of domains. We speculate that this observation suggests novel ways in which Homer may have evolved to regulate molecular decision-making at the post-synaptic density, and may have broader ramifications for the design of family-specific EVH1 inhibitors.

## Methods

### Protein Constructs and Purification (BLI Constructs)

EVH1 domains used for BLI experiments were cloned into a pMCSG7 vector for recombinant expression in *Escherichia coli* (Supplementary Table 4). The plasmid was transformed into Rosetta2(DE3) cells (Novagen) and grown in 1L Terrific Broth (TB) media with 100 µg/ml ampicillin. Cells were grown while shaking at 37 °C until an OD600 of 0.5-0.6, upon which the cultures were placed on ice for 20 min. Following this, protein expression was induced with 0.5 mM isopropyl β-D-1-thiogalactopyranoside (IPTG) and the induced cells were grown by shaking at 18 °C overnight. Cultures were harvested by centrifugation, and the resulting pellets were resuspended in Lysis Buffer (20 mM HEPES pH 7.6, 500 mM NaCl, 20 mM imidazole) at a concentration of 0.2 g/mL before being frozen at −80 °C.

The next day, the pellets were thawed and supplemented with 0.2 mM phenylmethylsulfonyl fluoride (PMSF) protease inhibitor. The cells were lysed by sonication, centrifuged, and the resulting supernatant was incubated with 2 mL Ni-NTA resin (washed in Lysis Buffer) while rotating at 4 °C. After 1 hour, the resin was washed 2x with 20 mL Lysis buffer. The protein was eluted from the Ni-NTA resin with 8 mL Elution Buffer (20 mM HEPES pH 7.6, 500 mM NaCl, 300 mM imidazole). TEV protease was added to the sample at an approximate concentration of 1 mg TEV:50 mg EVH1. The sample was dialyzed overnight at 4 °C in 1 L of TEV Buffer (50 mM HEPES pH 7.5, 300 mM NaCl, 0.5 mM EDTA, 2 mM DTT).

The completeness of the TEV reaction was assessed by sodium dodecyl-sulfate polyacrylamide gel electrophoresis (SDS–PAGE). Next, the sample was applied to 2 mL Ni-NTA resin and washed with 8 mL Lysis Buffer. The flow-through and wash were pooled, filtered, and loaded onto an S75 26/60 sizing column equilibrated in Gel Filtration buffer (20 mM HEPES pH 7.6, 150 mM NaCl, 1 mM DTT). Fractions were collected and their purity was assessed via SDS-PAGE. Pure samples were pooled and concentrated prior to being flash-frozen and stored at −80 °C.

### Protein Constructs and Purification (Surface Display Constructs)

EVH1 domains used for bacterial cell surface display and SPOT arrays were cloned into a pDW363 vector for recombinant expression in *E. coli* (Supplementary Table 4). The plasmid was transformed into Rosetta2(DE3) cells (Novagen) and grown in 1 L TB media with 100 µg/ml ampicillin and 0.05 mM D-(+)-biotin for in vivo biotinylation. Protein expression was carried out as above.

Pellets were thawed and supplemented with 0.2 mM PMSF prior to lysis by sonication. The lysed sample was centrifuged, and the resulting supernatant was incubated with 2 mL Ni-NTA resin (washed in Lysis Buffer) while rotating at 4°C. After 1 hour, the resin was washed 2x with 20 mL Lysis Buffer. The protein was eluted from the Ni-NTA resin with 8 mL of Elution Buffer (20 mM HEPES pH 7.6, 500 mM NaCl, 300 mM imidazole). The eluted sample was filtered and loaded onto a S75 26/60 sizing column equilibrated in Gel Filtration Buffer (20 mM HEPES pH 7.6, 150 mM NaCl, 1 mM DTT, 5% Glycerol). Fractions were collected and their purity assessed via SDS-PAGE. Pure samples were pooled and concentrated prior to being flash-frozen and stored at −80 °C.

### Protein Constructs and Purification (SUMO-Peptides)

Peptides were cloned as SUMO fusion constructs and inserted into a pDW363 vector for recombinant expression in *E. coli* (Supplementary Table 4). The construct was transformed into Rosetta2(DE3) cells (Novagen) and grown in Lysogeny Broth (LB) media supplemented with 100 µg/ml ampicillin and 0.05 mM D-(+)-biotin for in vivo biotinylation. The cells were grown while shaking at 37 °C until reaching an OD600 of 0.7, upon which protein expression was induced with 1 mM IPTG at 37°C for 5 hours. Cultures were harvested by centrifugation, and the resulting pellets were frozen overnight at −80 °C. The next day, the pellets were thawed and resuspended in 4 mL B-PER (Thermo Fischer)/g cell pellet and supplemented with 0.2 mM PMSF. Samples were incubated at room temperature for 15 minutes, spun down, and the supernatant was loaded into columns containing 0.4 mL Ni-NTA resin equilibrated in Wash Buffer (20 mM Tris pH 8.0, 500 mM NaCl, 20 mM imidazole). The columns were washed 3x with 1 mL Wash Buffer and then eluted from the column with 1.5 mL Elution Buffer (20 mM Tris, 50 0mM NaCl, 300 mM imidazole pH 8.0). Samples were flash-frozen and stored at −80 °C.

### Surface Display Constructs

Control peptides for bacterial surface display were cloned as C-terminal fusions to the surface protein eCPX using a vector designed by the Daugherty group.^39^ Peptides were flanked by an N-terminal FLAG tag and a C-terminal c-Myc tag for quantification of peptide expression on the bacterial surface (Supplementary Table 4). These plasmids were transformed into chemically competent MC1061 *E. Coli* cells for FACS analysis.

The T7-pep library, consisting of the same construct design detailed above, was transformed into electrocompetent MC1061 cells (Lucigen) to maximize transformation efficiency.^38^ 25 µL of cells were pre-chilled on ice for 10 min. These cells were mixed with plasmid and were pulsed in a 0.1cm cuvette at 1.8 kV, 600 Ohm, 10 µF using a Biorad electroporator. The cuvette was immediately rinsed twice with 1mL of Super Optimal Broth media with 20 mM glucose (SOC) that had been pre-warmed at 37 °C. A serial dilution of the transformed cells was plated on LB + chloramphenicol (Cm) plated to assess transformation efficiency. The cells were recovered at 37 °C for 1 hour and then combined with 40 mL LB + Cm + 0.2% glucose for overnight growth. Cells were re-transformed between rounds of enrichment to minimize the possibility of cells developing growth advantages associated with mutations in the chromosomal DNA.

### FACS Sample Preparation

The following protocol was adopted from Hwang et al.^33^ All PBS buffers were supplemented with additional NaCl to a final concentration of 300 mM. Cultures consisting of 5 mL LB + 25 µg/mL Cm + 0.2% glucose (w/v) were inoculated overnight (37 °C) with MC1061 cells encoding arabinose-inducible eCPX-peptide fusion constructs. The next morning, cells were pelleted and used to inoculate a fresh culture of LB + 25 µg/mL Cm (no glucose). These cultures were grown until an OD600 of 0.5-0.6 (37 °C). Upon reaching this density, eCPX expression was induced with 0.04% arabinose (w/v) for 1.5 hours. Following expression, the OD600 was measured again, and an appropriate volume of culture was centrifuged to obtain a pellet of 1 x 10^7^ cells/FACs analysis sample. In the case of T7-pep library sorting, a total of 1 x 10^8^ cells were pelleted. The pellets were washed with PBS pH 7.4 + 0.1% BSA and then incubated with anti-FLAG antibody conjugated to allophycocyanin (anti-FLAG:APC; PerkinElmer) (diluted 1:100 into PBS pH 7.4 + 0.1% BSA; 30 µL/1 x 10^7^ cells) for 30 min at 4 °C. Simultaneously, biotinylated monomeric EVH1 domain (1 µM) was pre-tetramerized by incubation with streptavidin-phycoerythrin (SA-PE; ThermoFisher) for 30 min. The cells were pelleted and washed with PBS pH 7.4 + 0.1% BSA to remove excess anti-FLAG:APC. Cells were pelleted, resuspended in PBS pH 7.4 + 1% BSA at 25µL buffer/1 x 10^7^ cells, and incubated with pre-tetramerized EVH1 domain at a 1:1 for a final volume of 50 µL/sample for 1 hour at 4 °C. Following this incubation, the cells were transferred to a pre-wet Millipore MultiScreen 0.22 µm hydrophilic low protein-binding plate and the buffer was removed by vacuum. The cells were then washed twice with 200 µL PBS + 0.1% BSA before being resuspended in 250 µL PBS + 0.1% BSA for FACS analysis or sorting.

### Flow Cytometry and FACS Analysis

Single-clone bacterial surface display analysis was carried out on a FACSCanto instrument (BD Biosciences) with samples prepared as described above. For each sample, 10,000 events were observed. FACS data were processed using the commercially available software FlowJo (FlowJo^TM^ Software for Mac, Version 10.8.1. Ashland, OR: Becton, Dickinson and Company; 2023). Cells were gated on (1) forward scatter height (FSC-H) vs. side scatter height (SSC-H) and (2) side scatter width (SSC-W) vs. side scatter height (SSC-H). Cells which made it through these two gates were analyzed for their APC and PE signals to quantify peptide expression and EVH1 binding, respectively.

Enrichment sorting of the T7-pep library was carried out on a FACSAria instrument (BD Biosciences). Collected cells were gated three times: (1) FSC-H vs. SSC-H (2) SSC-W vs. SSC-H (3) APC vs. PE. A no-peptide negative control and two positive control peptides of different affinities for Homer1 (CAP2, DBN1) were used to set the final collection gate. With each round of sorting, the PE voltage was adjusted to ensure a similar percentage of control cells fell within the gate. Samples, prepared as described above, were oversampled at least 10-fold relative to the library population. After the initial round of selection against the naïve T7-pep library, we performed a negative, PE-only sort and collected the non-binding population in order to eliminate sequences that were binding to PE. Subsequently, we performed 3 more rounds of positive selection using Homer1, for a total of 4 positive selections. The final round of sorting was performed twice (Sort5a/5b), but for the purpose of analysis, these populations were combined. Sorted cells were collected in SOC media and recovered at 37 °C for 1 hour. The culture was then combined with LB + 25 µg/mL chloramphenicol + 0.2% glucose (w/v) culture for overnight growth (37 °C) and then miniprepped (Qiagen).

### Illumina Amplicon Preparation

The plasmids from each round of enrichment sorting were prepared for single-lane Illumina deep sequencing as follows. Miniprepped plasmids from each round of enrichment (100 ng DNA/sample) were PCR amplified with a common forward primer to introduce an i5 Illumina anchor (FWD PCR) and a unique reverse primer to introduce the i7 Illumina anchor and a unique 6-nucleotide index for demultiplexing (REV INDEX X). A complete list of these primers and their sequences is provided in Supplementary Table 1. Twelve cycles of amplification were carried out with Q5 High Fidelity DNA Polymerase (NEB) using an annealing temperature of 68 °C and a 1 min extension at 65 °C. The success of this amplification was verified using a 1.5% agarose gel. Each PCR product was purified using a double-sided SPRI bead cleanup (AMPure XP-Beckman). First, a right-sided selection (0.56x) was used to remove large library fragments. The SPRI beads were resuspended in a solution of 2.5 M NaCl and PEG 8000 (20% w/v), mixed with the PCR products, and then incubated for 5 min at room temperature. Next, the beads were magnetically separated and the supernatant removed. This supernatant was then used to perform a left-sided selection (0.85x) to remove library fragments and adapter dimers. A fresh sample of SPRI beads was resuspended in PEG solution and then combined with the supernatant from the right-handed selection. The beads were then magnetically separated, the supernatant discarded and washed twice with 200 µL of pre-chilled 80% ethanol. Following this wash, the ethanol was removed and the beads were dried for 10 minutes at room temperature. The final sample was eluted from the beads by resuspension in 11 µL of 10 mM Tris pH 8.0 and their concentration measured via NanoDrop. The samples were pooled and submitted for sequencing using a NextSeq500 instrument with paired-end reads.

### Illumina Data Processing

Unless noted, an in-house script was used for the following data processing. Forward and reverse reads from Illumina sequencing were merged using the software BBMerge with the default overlap requirement of 12 base pairs and a quality score of 20.^82^ Read counts for each pool were collapsed by amino-acid sequence, and we applied an initial quality filter requiring a sequence length of 36 amino acids as well as an input library read count filter >5. The remaining 741,742 sequences were clustered to remove redundancy using CD-Hit, yielding a final count of 311,560 unique sequences.^83^ For each sequence, the raw read count was converted to a frequency (F_i_) by dividing by the total number of reads within the respective pool. Sequences that failed to enrich in the first selection against the naive input library were categorized as “non-enriching”. For sequences that persisted through the first two rounds of selection, an enrichment score was calculated by taking the area under the curve of the following transformation:

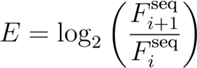

The distribution of this enrichment score was used to categorize sequences as “strongly enriching” or “weakly enriching” with a Z-score cutoff of 1.

### Sequence Logos

Matches to the core Homer motif were identified in sequences from the T7-Pep library using the regular expression [PAILV]P..[FWY]. Sequences were aligned by the motif with both an N- and C-terminal padding of 10 residues. In sequences where the motif was within 10 residues of the N-terminus, we appended the sequence “xxxxxxialr”, and for those within 10 residues of the C-terminus we appended “riarxxxxxx”. These residues correspond to non-proteomic sequences within the surface display construct and were displayed in lowercase to differentiate them from the proteomic residues within the logo. Position-specific scoring matrices (PSSMs) were generated using the publicly available software pLogo, with matches to the motif in all three populations (strongly enriching, weakly enriching, and non-enriching) serving as the background.^42^ PSSMs for sequences derived from the proteomic SPOT array were similarly aligned by the Homer motif. Final logos were generated from these PSSMs using the Logomaker python library.^84^

### BLI Assay

BLI experiments were carried out on an Octet Red96 instrument (ForteBio). All assay steps were carried out in a 1:1 mixture of EVH1 Gel Filtration Buffer (20 mM HEPES pH 7.6, 150 mM NaCl, 1 mM DTT) and PBS Buffer (1x PBS pH 7.6, 1 mM DTT, 1% BSA, 0.1% Tween-20). Data was collected at room temperature using a shake speed of 1000 rpm at the default sampling rate. Streptavidin-coated tips (ForteBio) were equilibrated in the above buffer for 10 min. Biotinylated SUMO-peptides were loaded on the streptavidin tips to a target optical density between 0.5-1 nm and subsequently washed in the buffer for 60 s. The loaded tips were immersed in a solution of EVH1 domain to collect association data for 100 s. Subsequently, EVH1-bound tips were transferred to a well containing buffer, and dissociation data was collected for 100 s. This process was done iteratively, using the same tip, with increasing concentrations of EVH1 domain, for a total of 8 concentrations.

Due to the fast kinetics of these interactions, we calculated K_D_ values using the equilibrium signal of the association step at each concentration of the EVH1 domain. This signal was calculated as the average signal over the final 10 seconds of association, minus the average of the final 10 seconds of the dissociation step. The association data of a SUMO-only control was subtracted from that of the SUMO-peptides. The binding curve was fit and K_D_ value was extracted in Kaleidagraph using non-linear least-squares fitting to the following equation:

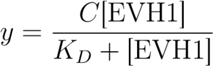

Where y is the signal and C is the max signal at saturation. K_D_ values and their respective errors are reported as the average and standard deviation on 3 technical replicates.

Fold change error in Figure 4C was calculated by error propagation using the formula:

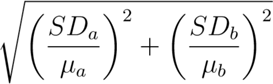

### Proteome Bioinformatic Analysis

Bioinformatic analysis of the human (UP000005640; Organism 9606) and *D. discoideum* (UP000002195; Organism 44689) proteomes was performed using the one protein per gene datasets available on UniProt. Regions were classified as disordered using IUPRED.^44^ Human orthologs for *D. discoideum* proteins were identified using the publicly available InParanoiDB9 (Human: 9606; Dictyostelium 44689).^85^

### SPOT Array

ABI3 substitution and overlapping motif SPOT arrays were synthesized at the MIT Biopolymer Facility using an Intavis SPOT synthesis peptide arrayer system. At the N-terminus, the 20-mer peptides were capped with a Gly-Ser linker and acetylated. The peptides were linked to the cellulose membrane by a C-terminal PEG spacer. For the overlapping motif array, because many of the peptide sequences contain additional Ena/VASP and Homer motifs, each peptide was also synthesized as a control where the internal proline tract was mutated to poly-Glycine. The sequence of each peptide is provided in Supplementary Table 2. The protocol for blocking and probing the membrane was adapted from Frank and Dubel.^86^ The membrane was hydrated in 100% ethanol, washed three times with TBS (50 mM Tris pH 7.0, 137 mM NaCl, 2.7 mM KCl, 1 mM DTT) and then blocked overnight in MBS (TBS +0.2% Tween-20, 2% (w/v) dry milk) at room temperature. The following day, the array was washed in T-TBS (TBS + 0.05% Tween-20) and then incubated with the protein sample in MBS (+1 µM DTT) for 3 hours at room temperature. All array experiments were carried out with streptavidin-Cy3 (ThermoFisher) pre-tetramerized EVH1 at a final monomer concentration of 0.5 µM. The membrane was washed three times in T-TBS prior to scanning on a GE Amersham Imager 680. SPOT intensities were calculated using the ImageJ analysis software.^87^ No background correction was performed. For the overlapping peptides, the final intensity was calculated as the difference between the intensity of the wild-type and motif knockout control. An intensity threshold of 1000 was used to classify a peptide as having bound, with peptides that exhibited an intensity >1000 for only one domain being annotated as selective. Overlapping motifs that bound to both ENAH and Homer1 via SPOT array were assessed as candidate binding partners based on localization (gTEX/UniProt subcellular localization), interaction (HURI/BioGRID), and GO term data (Figure S3).^43,88,89^ GO term analysis was performed using the SLiMSearch 4 “shared functional annotations” tool.^90^ A p-value threshold of 0.001 for any paralogous Ena/VASP or Homer family member was used as a cutoff for marking GO terms as being shared.

## Contributions

Avinoam Singer - Conceptualization, Data Curation, Formal analysis, Writing - Original Draft, Writing - Review and Editing. Alejandra Ramos - Data Curation. Amy E. Keating - Conceptualization, Funding Acquisition, Supervision, Writing - Original Draft, Writing - Review and Editing.

## Funding Information

Research reported in this publication was supported by the National Institute of General Medical Sciences of the National Institutes of Health under R01 GM129007 and R35 GM149227. Avinoam Singer received support from the National Institute of General Medical Sciences training award T32 GM007287. The content herein is solely the responsibility of the authors and does not represent the official views of any of the funding organizations.

## Acknowledgements

We thank the Koch Institute’s Robert A. Swanson (1969) Biotechnology Center for peptide array synthesis and flow cytometry expertise and services. We would also like to thank the MIT Biophysical Instrumentation Facility for access to BLI instrumentation. Lastly, we would like to thank the following individuals for their helpful discussion: Sebastian Swanson, Jackson Halpin, Theresa Hwang, Dia Ghose, Jennifer Kosmatka, and Foster Birnbaum. A special thank you to Robert T. Sauer and Joseph Davis for their instrumental feedback throughout the course of this work.

## Notes

### Competing Interest Statement

The authors have declared no competing interest.

### Summary of Updates

The revised manuscript makes minor changes to the tables, text, and funding information provided in the initial manuscript.

