## Supplementary material for "Elaboration of the Homer1 Recognition Landscape Reveals Incomplete Divergence of Paralogous EVH1 Domains": Figure S

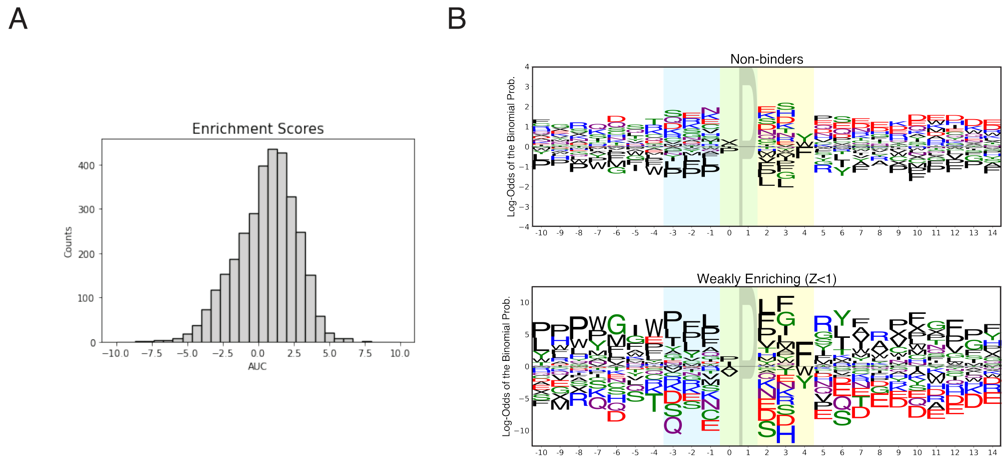


**Figure S1**: (A) Distribution of enrichment scores for T7-pep peptides. (B) Sequence logos for non-enriching and weakly enriching T7-pep peptides.

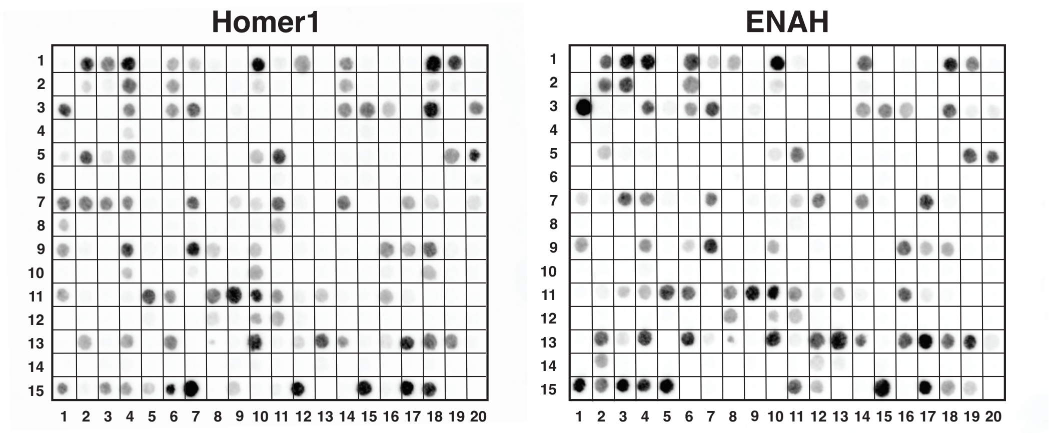


**Figure S2**: SPOT arrays for Homer1 and ENAH binding to human peptides containing overlapping Ena/VASP and Homer motifs. Corresponding sequences are provided in Supplementary Table 2.

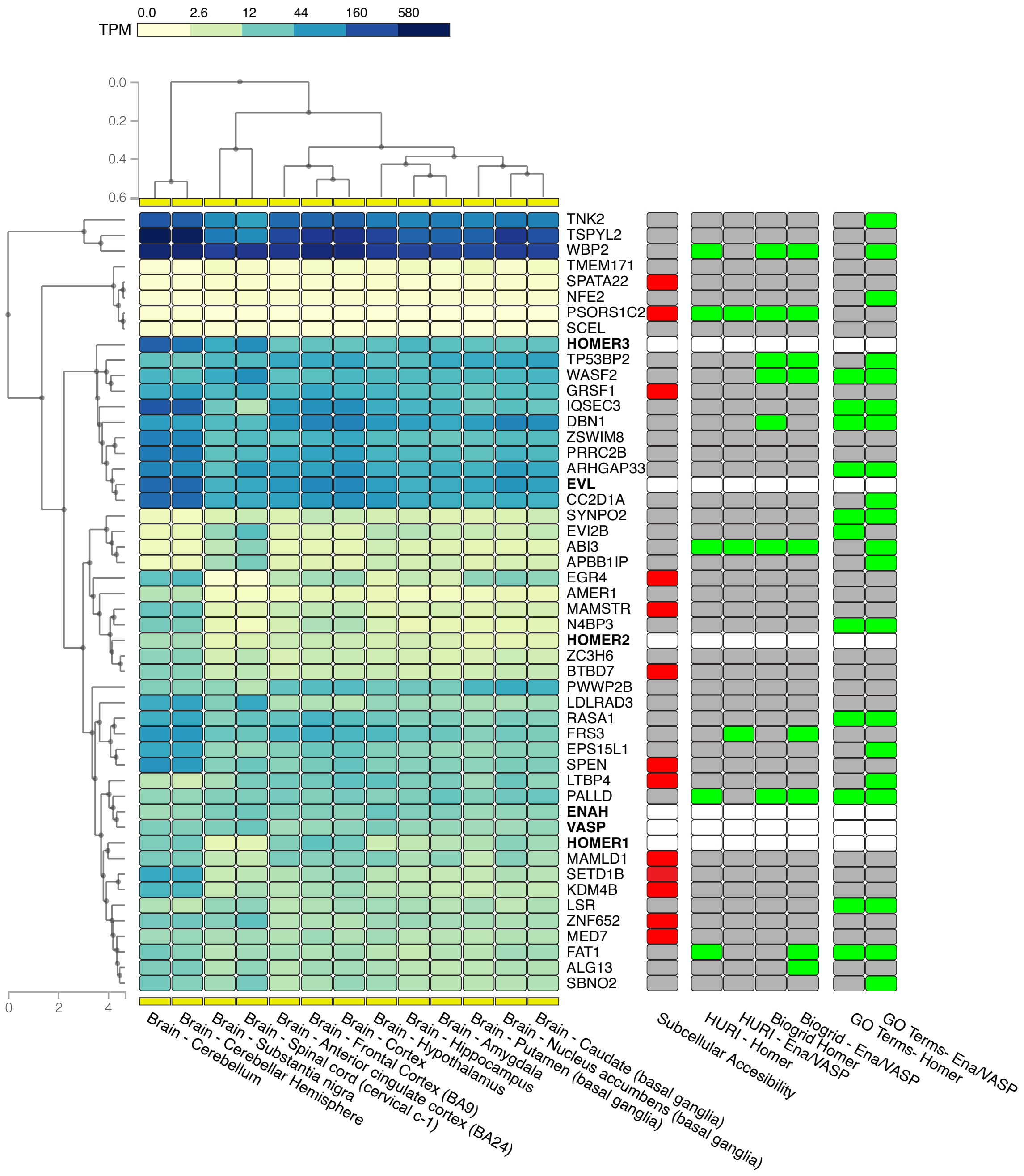


**Figure S3**: Expression, localization (gTex), and cataloged interactions of proteins containing promiscuous SLiMs (gTex).

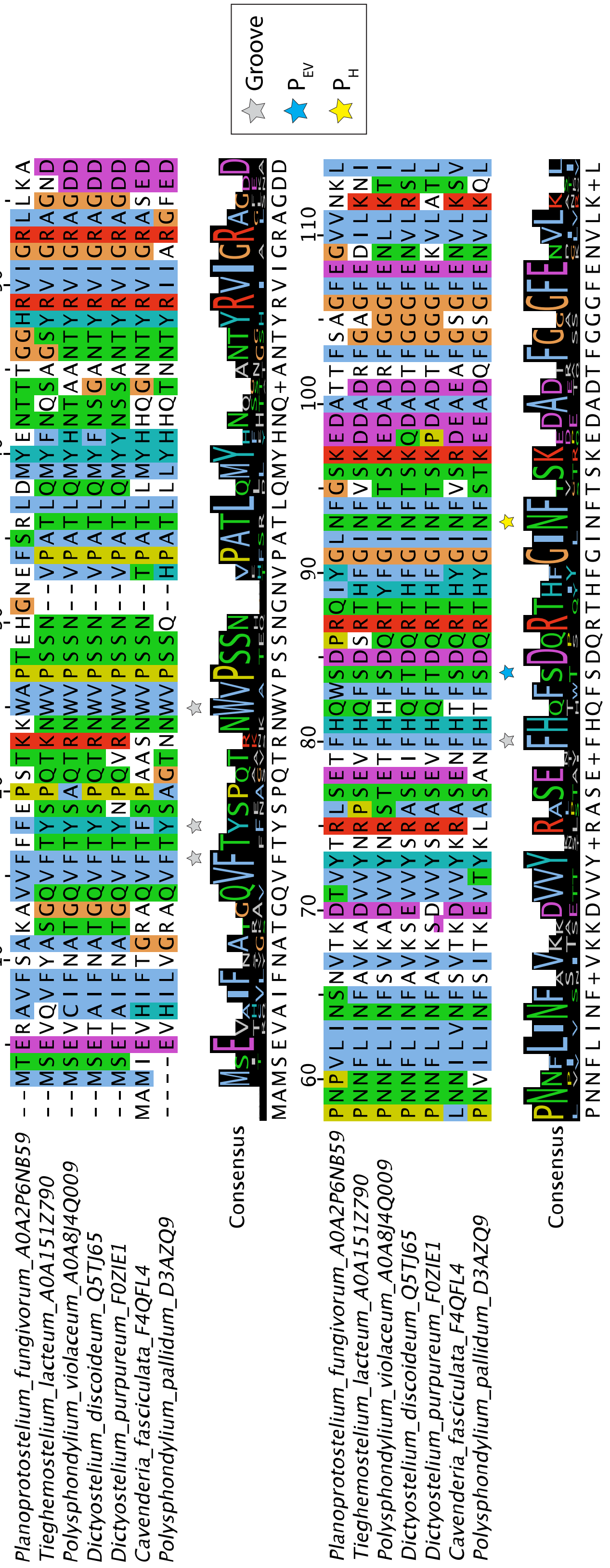

**Figure S4**: Sequence alignment of Ena/VASP EVH1 domains from amoebas.

A

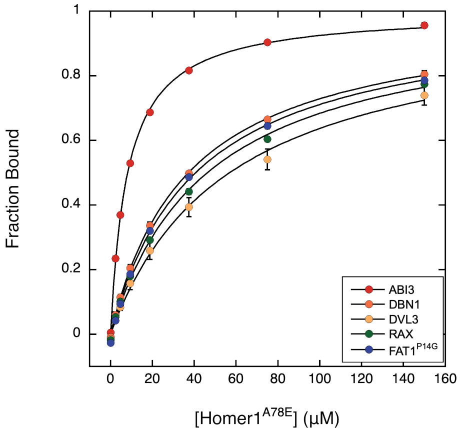


| **Name** | **Sequence** | **$K_D$ (µM)** |
|---|---|---|
| DBN1 | LNFDE**LPEPPATF**CDPEEVE | 37.3 ± 1.7 |
| FAT1$^{P5G}$ | DIESD**FPPPPEDF**GAADELP | 40.7 ± 1.4 |
| ABI3 | GDELG**LPPPPPGF**GPDEPSW | 8.2 ± 0.2 |
| RAX | GFGPPAQSLPASYTPPP**PPPPF**LNSPPLGPGLQPLA | 46.4 ± 1.4 |
| DVL3 | PLPHPGAAPWPMAFPYQ**YPPPPHPY**NPHPGFPELGY | 57.4 ± 8.0 |

B

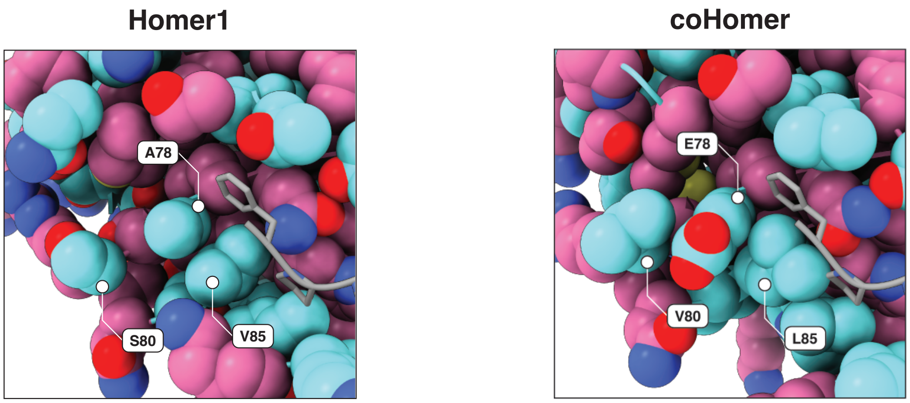


**Figure S5**: (A) BLI measurements for Homer$^{A78E}$. Values represent the average and standard deviation of 3 replicates. (B) Comparison of the structural context of E78$_{coHomer}$ (AlphaFold2) and A78$_{Homer1}$ (1DDV).
